# Dopamine modulates adaptive forgetting in medial prefrontal cortex

**DOI:** 10.1101/2021.04.08.438979

**Authors:** Francisco Tomás Gallo, María Belén Zanoni Saad, Juan Facundo Morici, Magdalena Miranda, Michael C. Anderson, Noelia V. Weisstaub, Pedro Bekinschtein

## Abstract

Active forgetting occurs in many species, but how the mechanisms that control behavior contribute to determining which memories are forgotten is still unknown. We previously found that when rats need to retrieve particular memories to guide exploration, it reduces later retention of other memories encoded in that environment. As with humans, this retrieval-induced forgetting relies on prefrontal control processes. The dopaminergic input to the prefrontal cortex is important for executive functions and cognitive flexibility. We found that, in a similar way, prefrontal dopamine signaling through D1 receptors is required for retrieval-induced forgetting in rats. Blockade of medial prefrontal cortex D1 receptors as animals encountered a familiar object impaired forgetting of the memory of a competing object in a subsequent long-term memory test. Inactivation of the ventral tegmental area produced the same pattern of behavior, a pattern that could be reversed by concomitant activation of prefrontal D1 receptors. We observed a bidirectional modulation of retrieval-induced forgetting by agonists and antagonists of D1 receptors in the medial prefrontal cortex. These findings establish the essential role of prefrontal dopamine in the active forgetting of competing memories, contributing to the shaping of retention in response to an organisms’ behavioral goals.

## Introduction

Much of what we experience is ultimately forgotten. Neuroscientific accounts of this inescapable process often have focused on the passive decay of memory traces (Davis and Zhong, 2017). However, recent neurobiological studies indicate that active forgetting mechanisms also can dictate a memory’s fate (Berry et al., 2012; Akers et al., 2014; Liu et al., 2016; Migues et al., 2016; Davis and Zhong, 2017; Awasthi et al., 2019). A common feature of both active forgetting processes and passive decay is that they are indifferent to memory content, but there is the question of whether forgetting of particular traces may be adaptively prioritized to benefit the organism’s goals. In human research on forgetting, however, selective forgetting mechanisms have been described that adaptively tune the accessibility of memories to organisms’ behavioral demands (Anderson, 2003). When people and rats retrieve a past event, other memories that compete with and hinder retrieval are more likely to be forgotten (Anderson et al., 1994). This ‘retrieval-induced forgetting’ occurs for a broad range of stimuli and contexts (Anderson and Hulbert, 2021; Anderson and Marsh, 2021). In humans, RIF arises because trying to retrieve a specific memory triggers inhibitory control mechanism mediated by the lateral prefrontal cortex that focus retrieval on goal-relevant traces by suppressing distracting memories (Anderson and Spellman, 1995; Anderson, 2003). Paralleling these findings rats can also engage this active forgetting mechanism to inhibit competing memories. As in humans, retrieval-induced forgetting in rats requires prefrontal engagement during the selective retrieval practice phase (Wu et al., 2014; Bekinschtein et al., 2018), and yields long-lasting forgetting that generalizes across multiple retrieval cues (Bekinschtein et al., 2018). Because memory systems throughout the animal kingdom confront the need to selectively retrieve goal-relevant memories, these findings suggest that the inhibitory control process that inhibits competing memories is conserved across mammalian species. In mammals generally, the prefrontal cortex facilitates flexible behavior (Miller and Cohen, 2001; Dalley et al., 2004; Ragozzino, 2007; Aron et al., 2014) via control mechanisms that suppress habitual responses that might otherwise dominate goal-directed action as well as been associated with attentional processes (Dalley et al., 2004; Aron et al., 2014) In rodents, the medial prefrontal cortex (mPFC) has been associated with attentional and inhibitory control processes (Miller and Cohen, 2001; Dalley et al., 2004; Ragozzino, 2007). We propose that the prefrontal cortex also suppresses competing memories, initiating a key signal that triggers active forgetting, tuning this process adaptively to an organism’s behavioral demands (Bekinschtein et al., 2018).

Decades of research have established the neurotransmitter dopamine as essential for cognitive control mechanisms mediated by the prefrontal cortex of humans, monkeys and rodents (Robbins, 2005). Dopamine in the mPFC modulates processes such as working memory (Sawaguchi and Goldman-Rakic, 1991; Zahrt et al., 1997; Granon et al., 2000; Vijayraghavan et al., 2007), attention, and behavioral flexibility (Ragozzino, 2002; Floresco, 2013). The rodent mPFC receives a dopaminergic input from neurons in the ventral tegmental area (VTA) that innervate both pyramidal cells and interneurons. In particular, D1 dopamine receptors (D1R) in the mPFC are critical for mediating dopamine effects on cognitive functioning (Floresco et al., 2006). Interestingly, an imaging genetics study in humans has linked genetic variation in prefrontal dopamine levels to differences in the engagement of lateral prefrontal cortex during selective retrieval and, correspondingly, to adaptive forgetting (Wimber et al., 2011). The data show a gene-dose-dependent influence of catechol-O-methyltransferase (COMT) Val108/158Met genotype on behavioral and brain activity indices of retrieval-induced forgetting, that increased linearly with Met allele load, suggesting a positive relationship between cortical dopamine availability and inhibitory control over competing memories. In the present study, we further tested the parallels in retrieval-induced forgetting across species by using our adaptation of the spontaneous object recognition paradigm (Bekinschtein et al., 2018). Specifically, we investigated whether dopamine-mediated control processes in the rodent prefrontal cortex contribute to adaptive forgetting of competing memories in our rodent model of retrieval-induced forgetting. We found that blockade of D1R receptors in mPFC of rats abolished retrieval-induced forgetting of object memories, that inactivation of VTA activity also impaired forgetting and that this impairment was reversed by concurrent activation of D1R receptors in mPFC. In addition, we show that dopaminergic modulation of adaptive forgetting is bidirectional, as activation of D1R receptors in mPFC significantly enhances retrieval-induced forgetting. Our results suggest that dopamine-dependent mechanisms of cognitive control over memory are conserved across species and are essential for adaptive forgetting in the mammalian brain.

## Results

To test whether control processes regulated by dopamine in the mPFC participate in adaptive forgetting, we studied how exploratory behavior in a rodent object recognition task was affected by manipulation of the dopaminergic system. Rats as well as many other species innately prefer novel objects to familiar ones and, in displaying this preference, reveal memory for the familiar object (Berlyne, 1950; Ennaceur and Delacour, 1988; Thompson et al., 1991; Winters et al., 2008; Blaser and Heyser, 2015; May et al., 2016). As in our previous study, we capitalized on this tethering of innate behavior and cognition to show that remembering a prior encounter with one object caused rats to forget other objects seen in the same setting (Bekinschtein et al., 2018). We modified the spontaneous object recognition procedure to include three phases equivalent to the ones present in human studies of retrieval-induced forgetting (Anderson et al., 1994; Ciranni and Shimamura, 1999; Maxcey and Woodman, 2014; Wimber et al., 2015): encoding, retrieval practice and test. Briefly the task is divided in three conditions. Each condition is divided in three phases. (Figure 1A) (see Material and Methods).

**Figure 1.**
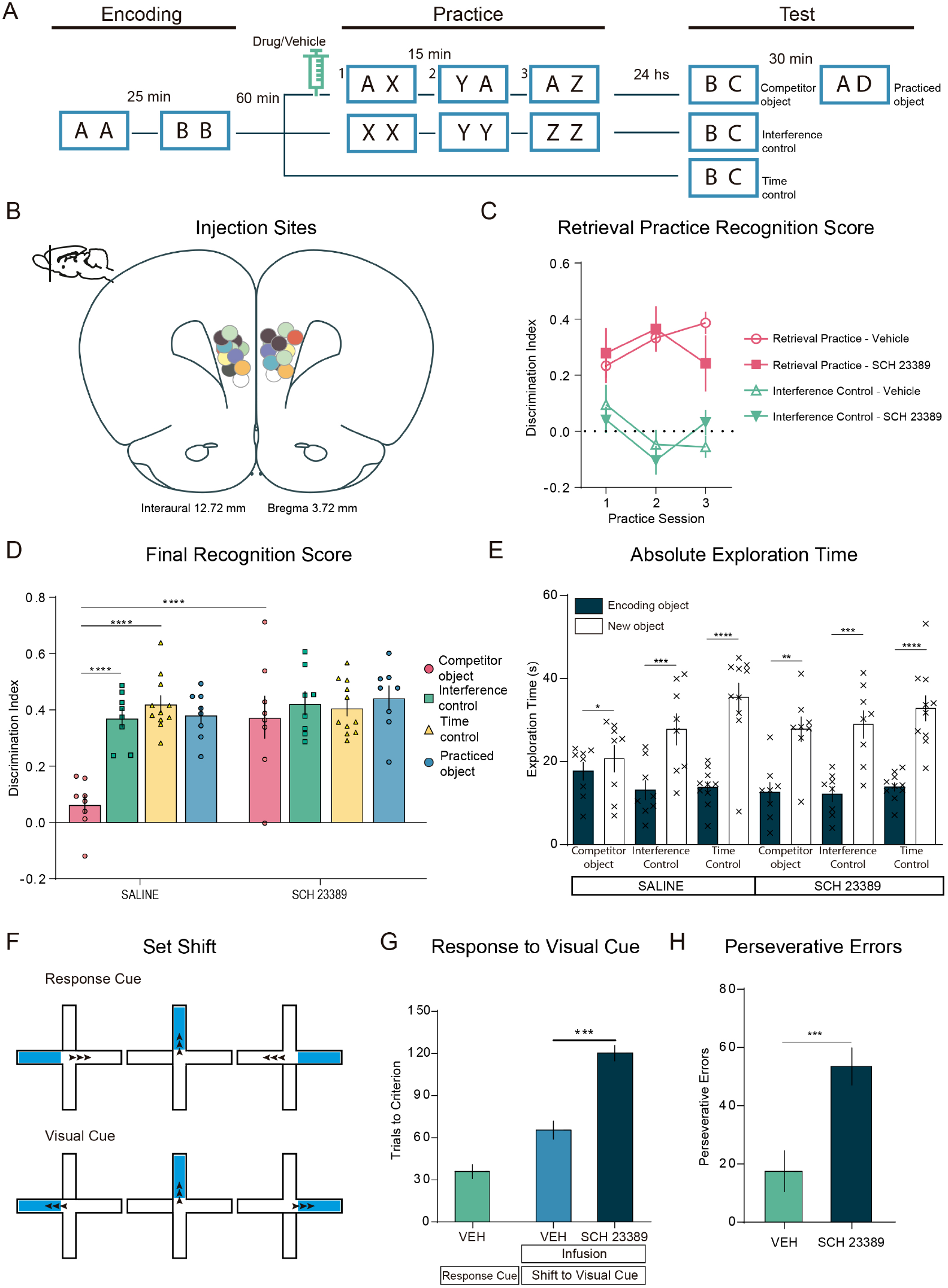
D1 receptors in medial prefrontal cortex mediate retrieval-induced forgetting. (A) Schematic representation of the behavioral protocol. After the acquisition, animals were divided into three different conditions, RP, IC and TC. The syringe indicates the infusion of the drug or its vehicle 10 minutes before the practice phase. **(B)** Histology. Diagram of the coronal section of the rat’s brain, showing the placement of the markings produced by methylene blue infusion for all the rats that received infusions of dopaminergic (or vehicle) drugs in the mPFC. The sections of the brain correspond to the atlas by Paxinos and Watson (1998)(Paxinos and Watson, 1998). **(C)** Discrimination indexes for the three sessions of the practice phase for the RP and IC groups in drug conditions and their vehicle (Table 1). **(D)** Discrimination indexes ± SEM for the testing phase. Animals did the task twice, once with the drug and once with the vehicle in a pseudorandomized way and for the same condition (Table 2), Two-way ANOVA, n=8-11, Bonferroni post hoc comparisons are shown indicated by asterisks. **(E)** Exploration times ± SEM for each individual object in the test phase (Table 2) compared by a Paired t test, shown with asterisks. **(F)** Training schemes for the set shifting task, with the Response Cue (left, egocentric) and the Visual Cue (right, visual). The arrows indicate the correct turn expected for each example trial. **(G)** Trials to criterion ± SEM is the number of trials conducted to complete a Criterion test correctly. Ordinary One-way ANOVA, n=5, Tukey post hoc comparisons are shown indicated by asterisks. **(H)** Perseverative Errors ± SEM, each trial in which the animal responded according to the self-centered key. Perseverative errors were defined as entering the wrong arm in three or more trials per block. Unpaired t test comparisons are shown by asterisks. *p<0.05, **p<0.01, ***p<0.001 and ****p<0.0001.

The D1 receptor (D1R) is one of the main dopamine receptors in the mPFC (Sawaguchi and Goldman-Rakic, 1991; Arnsten, 1998). Thus, in Experiment 1 we studied the role of mPFC D1R in retrieval-induced forgetting. Rats were implanted with cannulae reaching the mPFC before the beginning of the experiment and were tested twice in each condition (retrieval-practice (RP), interference control (IC) or time control (TC), once with saline and once with the D1R antagonist SCH23390 (SCH). We injected SCH (0.3 μg/μl, 0.5 μl per side) into the mPFC bilaterally (Figure 1B) 10 min before the first retrieval practice trial, and at the same point in the IC and TC conditions.

**Table 1:**
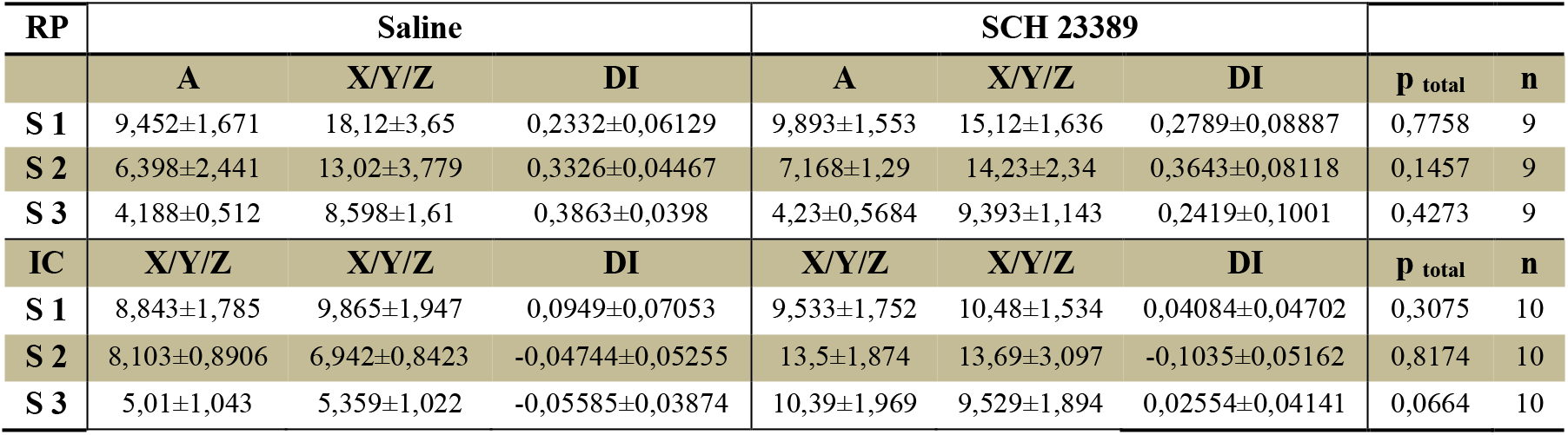
Exploration times and discrimination indexes during the practice phase in the Retrieval Practice condition for experiment depicted in Fig 1A. **Retrieval practice phase.** Total exploration times during the retrieval practice phase and DI for the RP and IC groups when animals were infused with saline (left) or SCH 23389 (right). Values are expressed in seconds (mean ± S.E.M.). Student’s t test, comparing DI between saline- and SCH-injected animals for each retrieval practice session (e.g. SCH 23389 A+X mean vs saline A+X mean for RP group and SCH 23389 X_1_+X_2_ mean vs saline X_1_+X_2_ mean, for IC group). Significance level is indicated as “p_total_”. SCH 23389 injection did not affect total exploration times during the practice phase compared to saline injection.

**Table 2:**
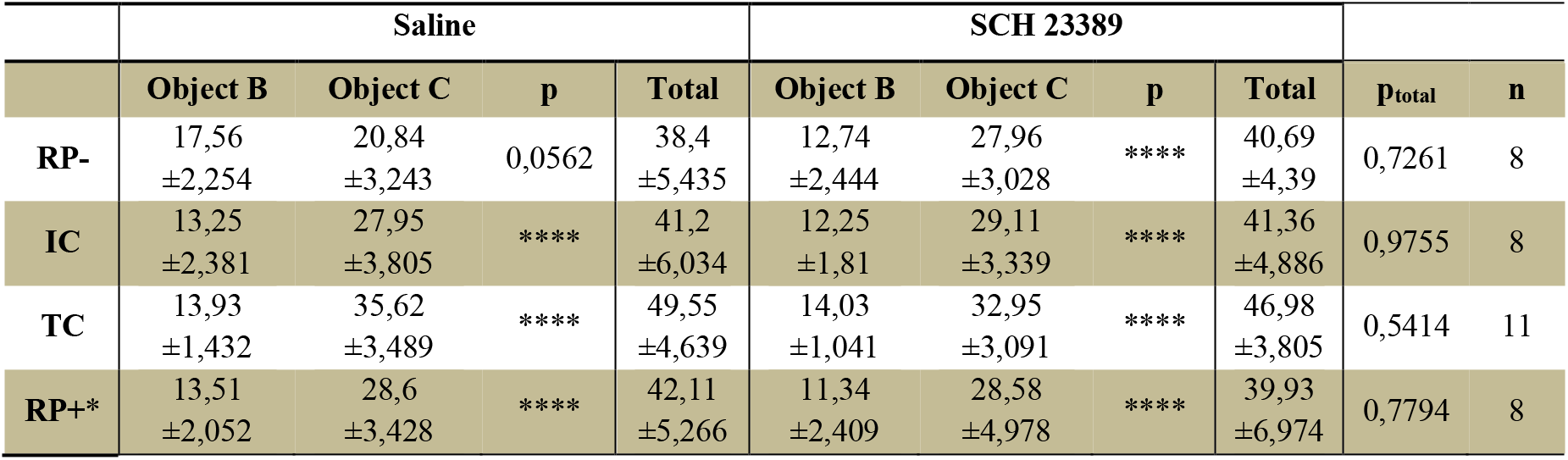
Exploration times during the final test phase for experiment depicted in Fig. 1A. **Absolute exploration times during the final test phase.** Total exploration times during the final test phase for the RP-, IC, TC and RP+ conditions. Values are expressed in seconds (mean ± S.E.M.). Paired student’s t test, comparing individual object exploration time between saline- and SCH-injected animals for the test phase; significance level is indicated as “p”. Paired student’s t test, comparing total exploration time between saline- and SCH-injected animals for the test phase (e.g SCH 23389 B+C mean vs saline B+C). Significance level is indicated as “p_total_”. *: RP+ group was exposed to the practiced object ‘A’. *: p<0,05; **: p<0,01; ***: p<0,001; ****: p<0,0001.

Although SCH injection still could have affected retrieval practice performance, we observed no evidence of this in any of the conditions: infusing animals with saline or SCH did not alter their total exploration times during this phase (total exploration times: RP veh: 51.12 s ±5.741; RP SCH: 59.21 s ±6.025, n=9, paired t test, p=0.2402, t= 1.269, df= 8). For the RP group, in both the saline and SCH conditions, rats preferred the novel objects during practice trials, indicating that retrieval of the practiced object was not affected by SCH infusion (Figure 1C, Table 1).

On the final test, we scored the time rats spent exploring the old object versus the novel object (Figure 1E). We computed a discrimination index reflecting the bias in the time they spent exploring the novel item instead of the old one (Figure 1D). We considered there was a retrieval-induced forgetting effect when the discrimination index of the RP group for the competitor object at test was significantly lower than the discrimination index of the IC and TC groups. We found that rats administered with saline showed evidence of intact retrieval-induced forgetting, as previously shown (Bekinschtein et al., 2018): saline-injected rats explored the competitor object B as if it was new as shown by the lower discrimination index in the RP condition compared with the IC and TC groups (Figure 1D, Table 2). Critically, however, rats injected with SCH showed impaired retrieval-induced forgetting (two-way ANOVA, Interaction: p= 0,0013, F (3, 31)= 6,654, Drug: p= 0,0008, F(1, 31)= 13,77, Condition: p< 0,0001, F(3, 31)= 10,05, Subjects: p= 0,3591, F(31, 31)= 1,140). Bonferroni corrected comparisons confirmed that rats’ memory for competitors was worse when injected with saline than with SCH. Indeed, injecting SCH abolished retrieval-induced forgetting completely (Figure 1D). The discrimination index in the RP group was indistinguishable from that of the IC or TC groups. Taken together, these findings support our hypothesis that inhibitory control mechanisms dependent on dopaminergic function in the mPFC are essential for retrieval-induced forgetting.

A different group of rats injected with SCH or Veh were evaluated in a set-shifting task that requires the organism to exert inhibitory control over the tendency to engage in a previously relevant behavioral strategy (Ragozzino et al., 1999; Birrell and Brown, 2000; Stefani et al., 2003). Blockade of D1R in mPFC has been shown to impair performance in this task (Ragozzino, 2002; Floresco et al., 2006). Set-shifting was conducted as in Floresco et al (2006)(Floresco et al., 2006), briefly there were 5-7 days of habituation to the maze, the handling and the food reward. The following days corresponded to the response discrimination training and the shift to visual cue learning. For the response discrimination training, the animal was required to always turn in one direction (opposite to its turn bias, left or right), regardless of the location of the visual cue placed in one of the arms. After the rat achieved acquisition criterion, it received a probe trial that consisted of starting the rat from the fourth arm that was not used as a start arm during testing. The day after reaching criterion on the response version, rats were now trained to enter the arm that contained the visual cue. Each rat was injected with SCH or Veh into the mPFC 15 min before the beginning of the visual cue learning session. The training procedure was similar to that used in the response version (see *Materials and Methods*). Errors were scored as entries into arms that did not contain the visual cue. As expected, blockade of D1R receptors in the mPFC impaired shifting from a response to a visual cue strategy. SCH-injected rats produced significantly more errors than Veh-injected animals in the probe trials and required significantly more of trials to reach criterion (Figure 2G, Table 3, Acquisition Criterion. One-way ANOVA: treatment: p< 0.0001, F(2, 17)= 85,92, Response vs. Visual Veh, p< 0.01, Response vs. Visual SCH, p< 0.0001, Visual Veh vs. SCH, p< 0.0001. Trials to Criterion. One-way ANOVA: treatment: p< 0.0001, F(2, 17)= 149,5, Response vs. Visual Veh, p< 0.0001, Response vs. Visual SCH, p< 0.0001, Visual Veh vs. SCH, p< 0.0001). Blocking D1R receptors in mPFC also affected the shift from an egocentric strategy to a visual strategy. Animals infused with SCH increased the number of trials to achieve the criterion relative to vehicle-infused animals (Figure 2G, Table 3, stats; Acquisition Criterion and Trials to Criterion) and made a greater number of perseverative errors (Figure 2H, Table 3, Unpaired t test: Perseverative Errors: p= 0,0173, t= 2,990, df= 8; Total Perseverative Errors: p< 0,0001, t= 9,856, df=8). Thus, blockade of D1R-dependent dopamine signaling with the same dose of SCH impaired both cognitive control and retrieval-induced forgetting.

**Table 3:** Set-Shifting parameters for the experiment depicted in Fig 1 F. **Set-shifting parameters.** Acquisition criterion, defined as the total number of test trials to complete 10 consecutive correct choices in a session. Trials to criterion, defined as the total number of test trials completed before a correct choice on the probe trial was made. Probe trials, defined as the total number of probe trials to get one correct. Perseveration involved continuing to make the same egocentric response, as required on the response version, when the trial required turning the opposite direction to enter the visual-cue arm. Perseveration was defined as entering the incorrect arm in 3 or more trials per block. After a rat stopped perseverating, the number of errors was counted when a rat reverted back to previously correct response (regressive errors) on those same type of trials that required the opposite turn as on the response version. Never reinforced errors were counted whenever a rat made an error by turning into the opposite response cue (with visual cue) arm.

|  | Response Cue | Visual Cue |  | One-way ANOVA | Response vs Visual |
| --- | --- | --- | --- | --- | --- |
|  |  | Saline | SCH 23389 |  |  |
| Acquisition Criterion | 34,15 ±3,656 | 55,67 ±4,616 | 109,7 ±3,593 | p<0.0001<br>F (2, 17) = 85,92 |  |
| Trials to Criterion | 20,31 ±3,718 | 65,33 ±4,072 | 118,5 ±3,801 | p<0.0001<br>F (2, 17) = 149,5 |  |
|  |  |  |  |  | Unpaired T Test |
| Total Perseverative Errors | - | 17,00 ±2,775 | 53,80 ±2,498 | - | p<0,0001<br>t=9,856 df=8 |
| Perseverative Errors | - | 5,600<br>±0,9798 | 35,20 ±6,888 | - | p=0,0173<br>t=2,990 df=8 |
| Regressive Errors | - | 12,60 ±2,542 | 18,40 ±5,972 | - | p= 0,3976<br>t=0,8936, df=8 |
| Never Reinforced Errors | - | 8,00 ±2,214 | 1,400<br>±0,5099 | - | p= 0,0196<br>t=2,910, df=8 |
| n | 10 | 5 | 5 |  |  |

**Figure 2.**
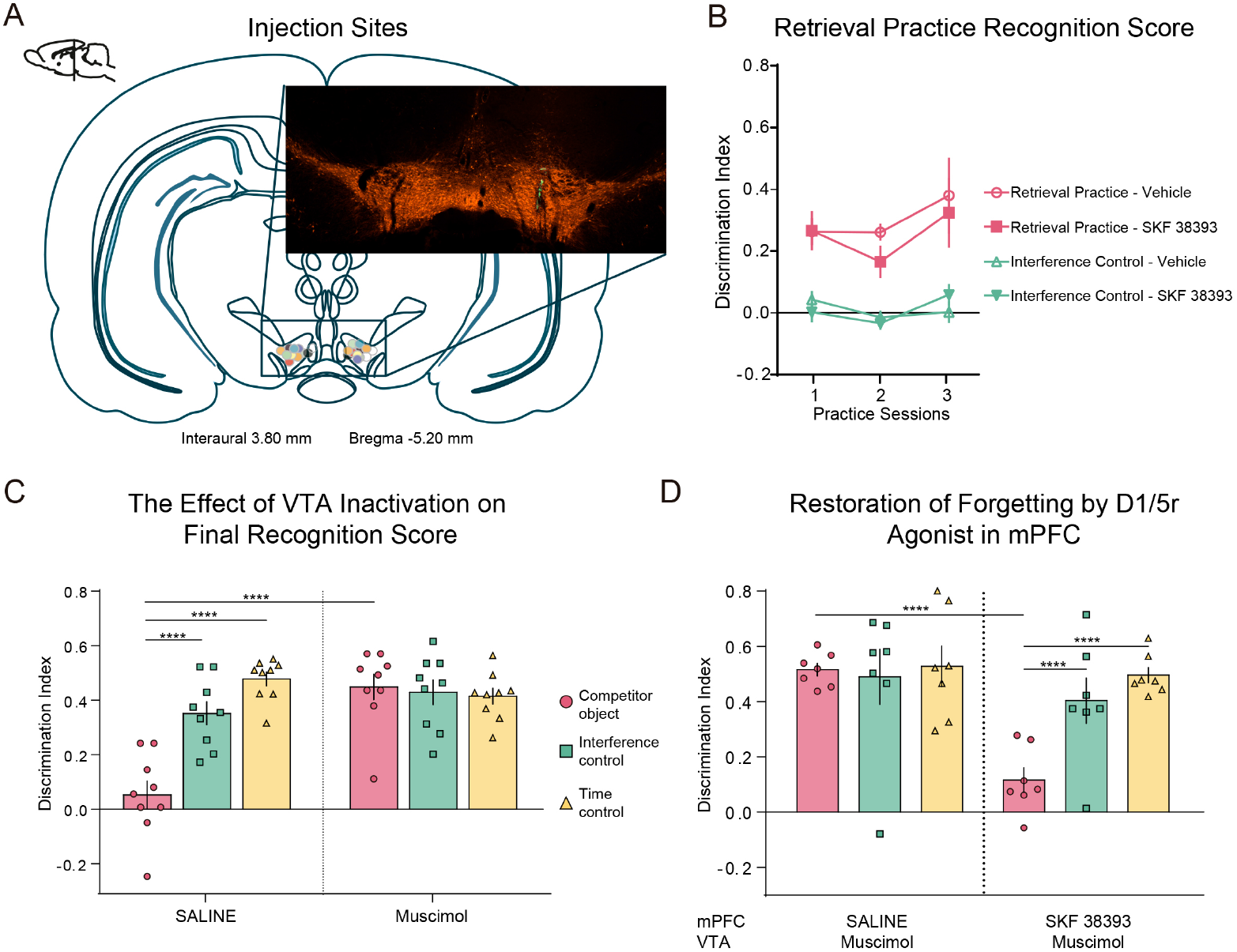
VTA projections to mPFC are necessary for retrieval-induced forgetting. **(A)** Diagram of coronal section of rat brain, showing the site of infusion of fluorescent green beads for all rats injected with muscimol (or vehicle) in the VTA. The sections of the brain correspond to the atlas by Paxinos and Watson (1998). Immunofluorescence, in orange anti-TH, in green, Green Beads infused through the implanted cannula. **(B).** Discrimination indexes ± SEM for the three sessions of the practice phase for the RP and IC groups under both conditions **(C)** Discrimination indexes ± SEM for the test phase after Mus or Veh injection into the VTA. Two-way ANOVA with Bonferroni’s post hoc analysis. There was a significant drug × condition interaction. Muscimol impaired the forgetting of the competitor object. **(D)** Discrimination indexes ± SEM for the test phase of the ‘Restoration of forgetting’ experiment by infusion of SKF38393 in mPFC. The animals did the task twice, once with the drug and once with the vehicle in a pseudorandomized way for the same condition. All animals were infused with muscimol in the VTA. Two-way ANOVA followed by a Bonferroni’s post hoc analysis indicated a significant drug × condition interaction. **p<0.01, ***p<0.001 and ****p<0.0001.

The main prefrontal dopamine source is the ventral tegmental area (VTA), which projects directly to the mPFC (Berger et al., 1991). We designed experiment 2 to establish whether dopamine release from VTA terminals into mPFC was required for retrieval-induced forgetting. We injected bilaterally the GABA agonist Muscimol (Mus, 0.1 mg/ml) or vehicle (Veh, saline) directly into VTA 15 min before the first retrieval practice trial (Figure 2A). Unlike permanent lesions, this treatment causes a transient silencing of the structure (Mao and Robinson, 1998) allowing the final memory test to occur in the absence of the drug. Injections were also made before exposure to the interpolated objects (equivalent to the “practice phase”) in the IC condition or before returning rats to their homecages for the TC condition.

Mus injection in VTA did not affect total object exploration during the practice phase (Total exploration times: RP veh: 92.99 s ±9.354 n=10; RP SCH: 85.97 s ±8.634, n=12, unpaired t test, p=0.5880, t= 0.5505, df= 20) (Table 4). Critically, during the test phase, in the Veh-injected animals the discrimination index was significantly lower for the RP condition compared with the IC and TC groups, whereas we did not observe any difference between the RP and the control groups in Mus-injected animals (Figure 2C; two-way ANOVA: Interaction: p< 0,0001, F(2, 48)= 16,29, Drug: p= 0,0002, F(1, 48)= 16,49, Condition: p< 0,0001, F(2, 48)= 11,95; Bonferroni *post hoc* multiple comparisons) and exploration times (Table 5). Given that Mus had no effect in the IC or TC conditions (Figure 2C, Table 5), this indicates that silencing the VTA did not modify recognition memory, but rather that VTA activity during the practice phase was specifically required for successful forgetting of the competing object memory. These findings are consistent with our hypothesis that dopamine release into mPFC during selective retrieval practice was important for successful control processes that inhibited competing memories and produced retrieval-induced forgetting.

**Table 4:** Exploration times and discrimination indexes during the practice phase in the Retrieval Practice condition for experiment depicted in Fig 2 (muscimol into the VTA) **Retrieval practice phase.** Total exploration times during the retrieval practice phase and DI for the RP and IC groups when animals were infused with saline (left) or Muscimol (right). Values are expressed in seconds (mean ± S.E.M.). Unpaired Student’s t test, comparing total exploration time between saline- and SCH-injected animals for each retrieval practice session (e.g. Muscimol A+X mean vs saline A+X mean for RP group and Muscimol X_1_+X_2_ mean vs saline X_1_+X_2_ mean, for IC group). Significance level is indicated as “p_total_”. Muscimol injection did not affect total exploration times during the practice phase compared to saline injection.

| RP | Saline |  |  | Muscimol |  |  | p total | n |
| --- | --- | --- | --- | --- | --- | --- | --- | --- |
|  | A | X/Y/Z | DI | A | X/Y/Z | DI |  |  |
| S 1 | 11,31±1,426 | 20,92±4,216 | 0,2628±0,04375 | 10,76±1,561 | 22,04±4,753 | 0,2657±0,06394 | 0,8080 | 9 |
| S 2 | 10,11±0,8196 | 17,08±1,035 | 0,2611±0,02736 | 11,97±1,863 | 15,54±2,106 | 0,1652±0,05271 | 0,5380 | 9 |
| S 3 | 8,91±0,8228 | 24,95±3,23 | 0,3797±0,1228 | 7,692±1,447 | 17,86±3,33 | 0,3242±0,1127 | 0,3195 | 9 |
| IC | X/Y/Z | X/Y/Z | DI | X/Y/Z | X/Y/Z | DI | p total | n |
| S 1 | 9,825±1,236 | 10,85±1,367 | 0,04285±0,02872 | 12,84±2,024 | 12,6±2,305 | 0,002149±0,03356 | 0,3509 | 10 |
| S 2 | 12,46±0,7411 | 12,79±0,8129 | -0,0145±0,02118 | 15,75±2,983 | 16,81±3,137 | -0,03421±0,02047 | 0,2614 | 10 |
| S 3 | 12,89±2,218 | 13,15±2,03 | 0,002358±0,03524 | 12,98±1,508 | 14,85±2,005 | 0,05898±0,03443 | 0,7414 | 10 |

**Table 5:** Exploration times during the final test phase for experiment depicted in Fig 2, (muscimol into the VTA). **Absolute exploration time during the final test phase.** Total exploration times during the final test phase for the RP-, IC and TC conditions. Values are expressed in seconds (mean ± S.E.M.). Unpaired student’s t test, comparing individual object exploration time between saline- and Muscimol-injected animals for the test phase; significance level is indicated as “p”. Unpaired student’s t test, comparing total exploration time between saline- and Muscimol-injected animals for the test phase (e.g Muscimol B+C mean vs saline B+C mean). Significance level is indicated as “p_total_”. *: p<0,05; **: p<0,01; ***: p<0,001; ****: p<0,0001.

|  | Saline |  |  |  | Muscimol |  |  |  |  |  |
| --- | --- | --- | --- | --- | --- | --- | --- | --- | --- | --- |
|  | Object B | Object C | p | Total | Object B | Object C | p | Total | p<sub>total</sub> | n |
| RP- | 23,85<br>$\pm 1,928$ | 26,64<br>$\pm 3,237$ | 0,0411 | 49,49<br>$\pm 3,554$ | 15,03<br>$\pm 1,826$ | 39,73<br>$\pm 2,953$ | ** | 54,76<br>$\pm 3,84$ | 0,3294 | 9 |
| IC | 14,74<br>$\pm 1,919$ | 27,62<br>$\pm 4,587$ | *** | 42,36<br>$\pm 5,876$ | 15,58<br>$\pm 2,695$ | 31,95<br>$\pm 3,277$ | *** | 47,53<br>$\pm 4,438$ | 0,4931 | 10 |
| TC | 13,35<br>$\pm 2,321$ | 34,26<br>$\pm 5,904$ | **** | 47,61<br>$\pm 7,978$ | 13,83<br>$\pm 1,342$ | 29,9<br>$\pm 2,857$ | *** | 43,72<br>$\pm 3,601$ | 0,6629 | 8 |

In experiment 3 we sought to elucidate whether VTA projections to the mPFC where important to modulate activity in this structure and cause retrieval-induced forgetting, we combined Mus injections into the VTA with injection of the D1R agonist SKF38393 into the mPFC in a new set of animals. We reasoned that if activation of D1R in the mPFC by dopamine released from VTA terminals was a necessary step towards retrieval-induced forgetting, exogenous activation of D1R should reverse the effects of silencing the VTA. Mus was injected bilaterally into the VTA in all animals 15 min before retrieval practice (or the equivalent phase in the IC and TC conditions). Injection of SKF38393 (8.4 μg/μl, 0.5 μl per side) or Veh into the mPFC was performed 10 min before retrieval practice (or the equivalent phase in the IC and TC conditions). SKF38393 injection did not produce any changes in exploration or recognition of the familiar object during the practice phase (Total exploration times: RP veh: 52.17 s ±8.506; RP SKF 38393: 48.89 s ±4.141, unpaired t test, p=0.7350, t= 0.3464, df= 12; Table 6). Critically, SKF38393 administration into mPFC caused significant memory impairment in the RP group in the final test, compared with Veh-injected animals. Thus, SKF38393 completely reversed the effect of silencing VTA with Mus (Fig 2D; two-way ANOVA: Interaction: p= 0,0167, F(2, 36)= 4,598, Drug: p= 0,0022, F(1, 36)= 10,84, Condition: p= 0,0150, F(2, 36)= 4,729. Bonferroni *post hoc* multiple comparisons) and exploration times at test (Table 7). No differences in discrimination indexes were found between Veh and SKF38393-injected animals in the IC and TC groups (Figure 2D; Table 7). Thus, in the absence of activity within the VTA, activation of mPFC D1R was sufficient to produce retrieval-induced forgetting, indicating that activation of mPFC D1R via dopamine release from VTA is one of the main mechanisms required for retrieval-induced forgetting in rats.

**Table 6:** Exploration times and discrimination indexes during the practice phase in the Retrieval Practice condition for experiment depicted in Fig 2, (muscimol into the VTA and SKF 38393 into the mPFC). **Retrieval practice phase.** Total exploration times during the retrieval practice phase and DI for the RP and IC groups when animals were infused with Muscimol in the VTA and saline (left) or SKF 38393 (right) in the mPFC. Values are expressed in seconds (mean ± S.E.M.). Unpaired Student’s t test, comparing total exploration time between saline- and SCH-injected animals for each retrieval practice session (e.g. Muscimol A+X mean vs saline A+X mean for RP group and Muscimol X_1_+X_2_ mean vs saline X_1_+X_2_ mean, for IC group). Significance level is indicated as “p_total_”. Muscimol injection did not affect total exploration times during the practice phase compared to saline injection.

| RP | Muscimol VTA – Saline mPFC |  |  | Muscimol VTA – SKF 38393 |  |  |  |  |
| --- | --- | --- | --- | --- | --- | --- | --- | --- |
|  | A | X/Y/Z | DI | A | X/Y/Z | DI | p<sub>total</sub> | n |
| S 1 | 7,571<br>$\pm 1,207$ | 13,22<br>$\pm 2,516$ | 0,2278 $\pm 0,1363$ | 7,894<br>$\pm 0,6434$ | 14,18 $\pm 1,72$ | 0,2611<br>$\pm 0,0801$ | 0,7408 | 7 |
| S 2 | 5,286<br>$\pm 1,564$ | 10,28<br>$\pm 2,123$ | 0,3547 $\pm 0,1077$ | 4,904<br>$\pm 0,8004$ | 8,811<br>$\pm 1,656$ | 0,2395<br>$\pm 0,08151$ | 0,5012 | 7 |
| S 3 | 4,789<br>$\pm 1,339$ | 6,72 $\pm 1,429$ | 0,2163 $\pm 0,1641$ | 5,261<br>$\pm 2,345$ | 7,84 $\pm 1,751$ | 0,3204<br>$\pm 0,2087$ | 0,6835 | 7 |
| IC | X/Y/Z | X/Y/Z | DI | X/Y/Z | X/Y/Z | DI | p<sub>total</sub> | n |
| S 1 | 10,69<br>$\pm 1,366$ | 11,12<br>$\pm 1,519$ | 0,02109<br>$\pm 0,04114$ | 9,463<br>$\pm 2,341$ | 9,627<br>$\pm 2,147$ | 0,06022<br>$\pm 0,09299$ | 0,4248 | 7 |
| S 2 | 6,699<br>$\pm 1,045$ | 7,146<br>$\pm 1,005$ | 0,03768<br>$\pm 0,05354$ | 6,291<br>$\pm 0,7672$ | 7,533<br>$\pm 1,547$ | 0,04854<br>$\pm 0,1024$ | 0,8892 | 7 |
| S 3 | 5,166<br>$\pm 0,7348$ | 4,976<br>$\pm 0,6849$ | -0,01615<br>$\pm 0,04094$ | 3,569<br>$\pm 0,4883$ | 4,499<br>$\pm 0,6463$ | 0,1156<br>$\pm 0,03372$ | 0,5892 | 7 |

**Table 7:**
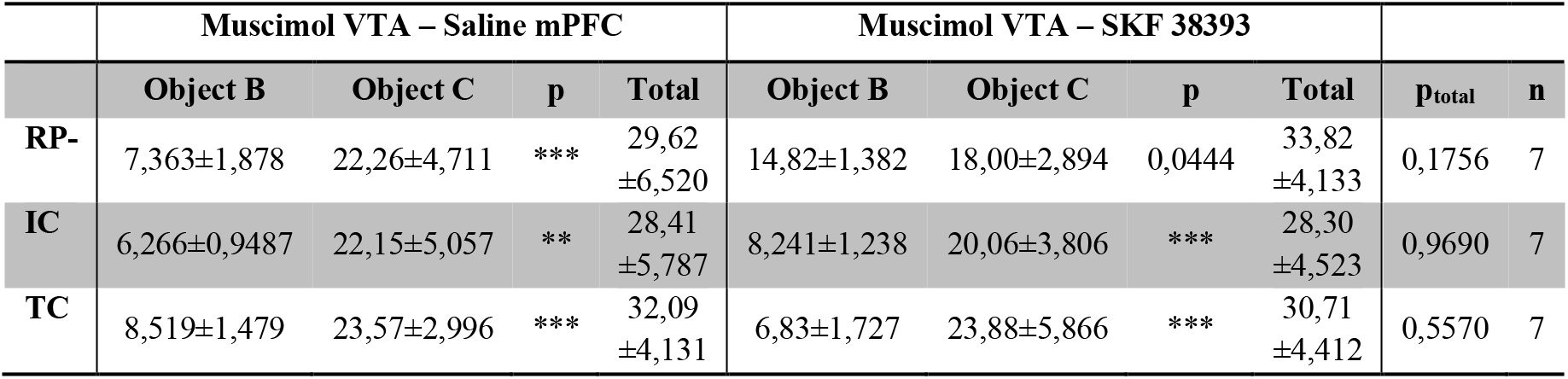
Exploration times during the final test phase for experiment depicted in Fig 2, (muscimol into the VTA and SKF 38393 into the mPFC). **Absolute exploration time during the final test phase.** Total exploration times during the final test phase for the RP-, IC and TC conditions. Values are expressed in seconds (mean ± S.E.M.). Unpaired student’s t test, comparing individual object exploration time between saline- and SKF-injected animals for the test phase; significance level is indicated as “p”. Unpaired student’s t test, comparing total exploration time between saline- and SKF-injected animals for the test phase (e.g SKF 38393 B+C mean vs saline B+C mean). Significance level is indicated as “p_total_”. *: p<0,05; **: p<0,01; ***: p<0,001; ****: p<0,0001.

In humans, higher prefrontal dopamine availability has been associated with greater retrieval-induced forgetting (Wimber et al., 2011). In rodents, D1R agonists have been shown to improve mPFC-related processes such as performance in a delayed-response task (Sawaguchi, 2001). It has been proposed that this enhancement could be due to diminished activity related to distracting information and/or increasing the signal-to-noise ratio (Williams and Goldman-Rakic, 1995; Vijayraghavan et al., 2007). If D1R activation in mPFC during the practice phase is required to maintain neural activity related to the target memory while minimizing the activity related to the distractor memory, then activation of these receptors could improve retrieval-induced forgetting. To evaluate this prediction, we injected the D1R agonist SKF38393 into mPFC in a new group of animals before a modified retrieval practice phase consisting of only one practice trial (Figure 3A). We reasoned that whereas only one practice trial would likely be insufficient to produce retrieval-induced forgetting on its own, it might do so given activation of D1R in mPFC, which could magnify the impact of inhibitory processes. A single retrieval practice did not yield significant memory impairment during the later test phase either in the Veh- or SKF-injected animals (Figure 3A, Table 9; two-way ANOVA: Interaction: p< 0,9014, F(4, 28)= 0,2594, Drug: p= 0,7148, F(1, 28)= 0,1363, Condition: p< 0,0001, F(4, 28)= 10,06. Bonferroni *post hoc* multiple comparisons). The impact of inhibition arising from one practice trial may have not been strong enough to produce retrieval-induced forgetting. In prior work, we had already observed that exposure to two retrieval practice trials during the practice phase induced RIF that was measurable in a test session 30 min after the practice phase (Bekinschtein et al., 2018). However, in the present study, the final test took place 24 h after the practice phase. Thus, we tested our hypothesis again, but with a protocol in which the animals were exposed to two practice trials as in our prior work (Bekinschtein et al., 2018) and injected with Veh or SKF (Figure 3A). In this case, we found no differences between Veh- or SKF-injected animals in the amount of RIF observed on the final test, as both groups showed similar and significant levels of retrieval-induced forgetting (Figure 3B, Table 11). Decreasing the number of trials proved not to be a sensitive strategy to evaluate positive modulation of retrieval-induced forgetting. We found an alternative approach to potentially observe a positive modulation of retrieval-induced forgetting. We introduced a longer delay in between the encoding phase, the practice phase and the final test, a manipulation that significantly reduced the size of retrieval-induced forgetting. We extended the delay between the encoding and final test phase to 48 h and the delay between the encoding and the retrieval practice phases to 24 h (see scheme in Figure 3C), with the aim of weakening the overall effect so that positive modulation could be observed. To ensure that memory performance was adequate to measure retrieval-induced forgetting after 48 hours, we modified our encoding protocol to create stronger memories. Preliminary work indicated that control animals required two separate exposures to each pair of objects during encoding to remember these objects 48 h later. Thus, we slightly modified the protocol for the particular mnemonic demands of longer lasting object memories.

**Table 8:** Exploration times and discrimination indexes during the practice phase in the Retrieval Practice condition for experiment depicted in Fig 3 A, (single practice session). **Absolute exploration times during the retrieval practice phase.** Total exploration times during the retrieval practice phase and DI for the RP-1 and IC1 groups when animals were infused with SKF 38393 or saline in the mPFC. Values are expressed in seconds (mean ± S.E.M.). Paired Student’s t test, comparing total exploration time between saline- and SKF-injected animals for each retrieval practice session (e.g. Saline A+X mean vs saline A+X mean for RP group and Muscimol X_1_+X_2_ mean vs saline X_1_+X_2_ mean, for IC group). Significance level is indicated as “p_total_”. SKF 38393 injection did not affect total exploration times during the practice phase compared to saline injection.

| RP-1 | Saline |  |  | SKF 38393 |  |  |  |  |
| --- | --- | --- | --- | --- | --- | --- | --- | --- |
|  | A | X/Y/Z | DI | A | X/Y/Z | DI | p total | n |
| S 1 | 11,42±1,668 | 17,53±2,202 | 0,2266±0,03074 | 16,25±1,858 | 23,15±3,381 | 0,1656±0,05351 | 0,1801 | 8 |
| IC1 | X/Y/Z | X/Y/Z | DI | X/Y/Z | X/Y/Z | DI | p total | n |
| S 1 | 18,03±1,241 | 17,05±1,744 | -0,03648±0,02197 | 18,44±2,371 | 19,28±2,586 | 0,02287±0,02644 | 0,2239 | 8 |

**Table 9:**
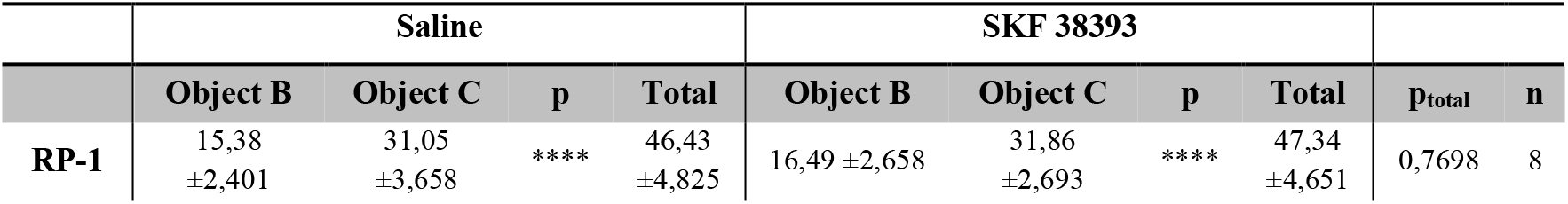
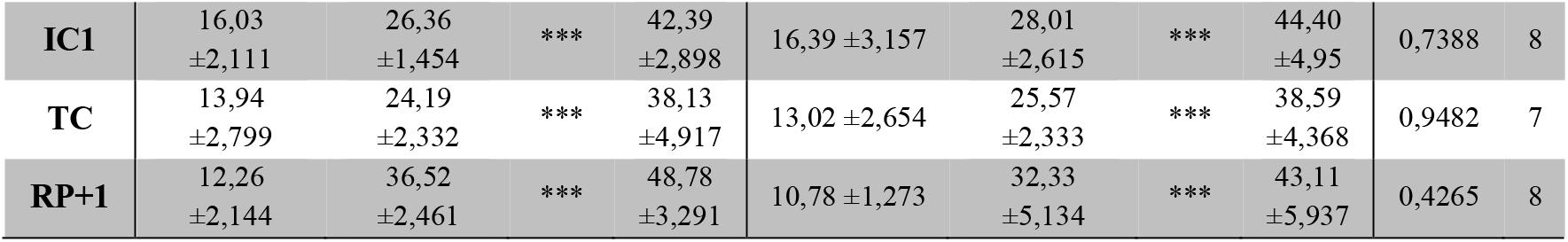
Exploration times and discrimination indexes during the final test phase for experiment depicted in Fig 3 A, (single practice session). **Absolute exploration time during the final test phase.** Total exploration times during the final test phase for the RP-1, IC1 and TC conditions. Values are expressed in seconds (mean ± S.E.M.). Paired student’s t test, comparing individual object exploration time between saline- and SKF-injected animals for the test phase; significance level is indicated as “p”. Paired student’s t test, comparing total exploration time between saline- and SKF-injected animals for the test phase (e.g SKF 38393 B+C mean vs saline B+C mean). Significance level is indicated as “p_total_”. *: p<0,05; **: p<0,01; ***: p<0,001; ****: p<0,0001.

**Table 10:**
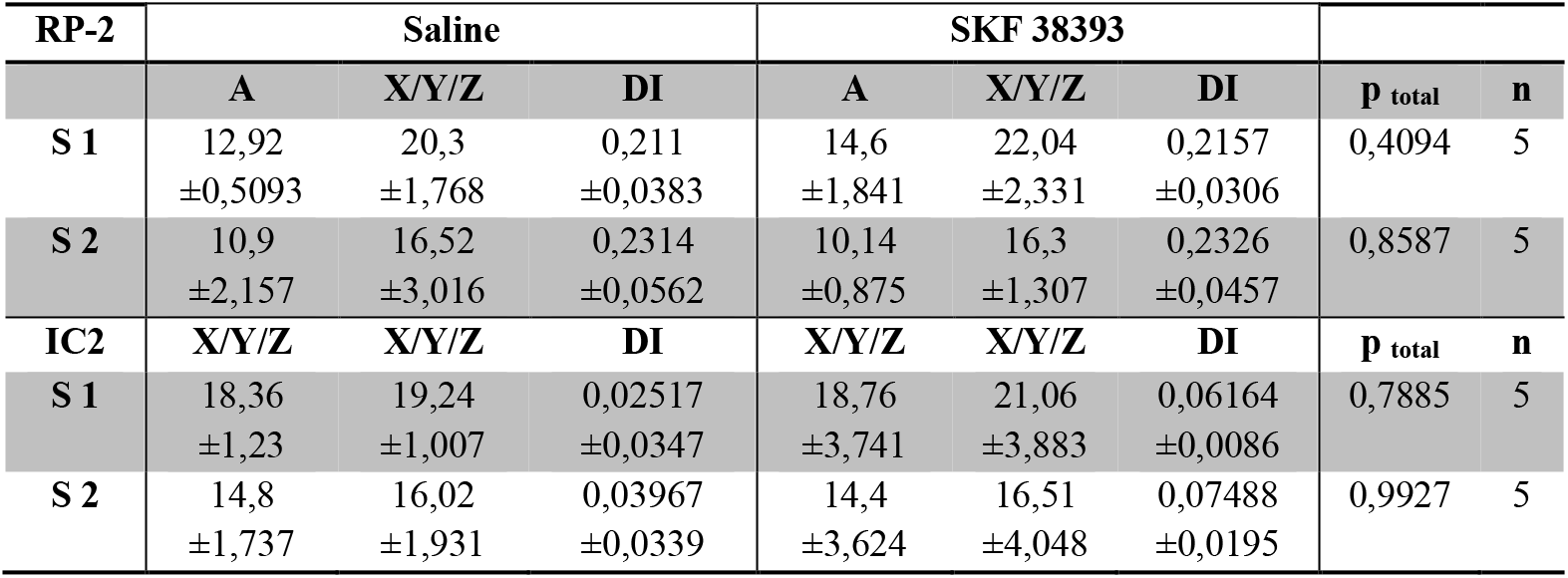
Exploration times and discrimination indexes during the practice phase in the Retrieval Practice condition for experiment depicted in Fig 3 A, (two practice sessions). **Retrieval practice phase.** Total exploration times during the retrieval practice phase and DI for the RP-2 and IC2 groups when animals were infused with SKF 38393 or saline in the mPFC. Values are expressed in seconds (mean ± S.E.M.). Paired Student’s t test, comparing total exploration time between saline- and SKF-injected animals for each retrieval practice session (e.g. Saline A+X mean vs SKF A+X mean for RP group and SKF X_1_+X_2_ mean vs saline X_1_+X_2_ mean, for IC group). Significance level is indicated as “p_total_”. SKF 38393 injection did not affect total exploration times during the practice phase compared to saline injection.

**Table 11:**
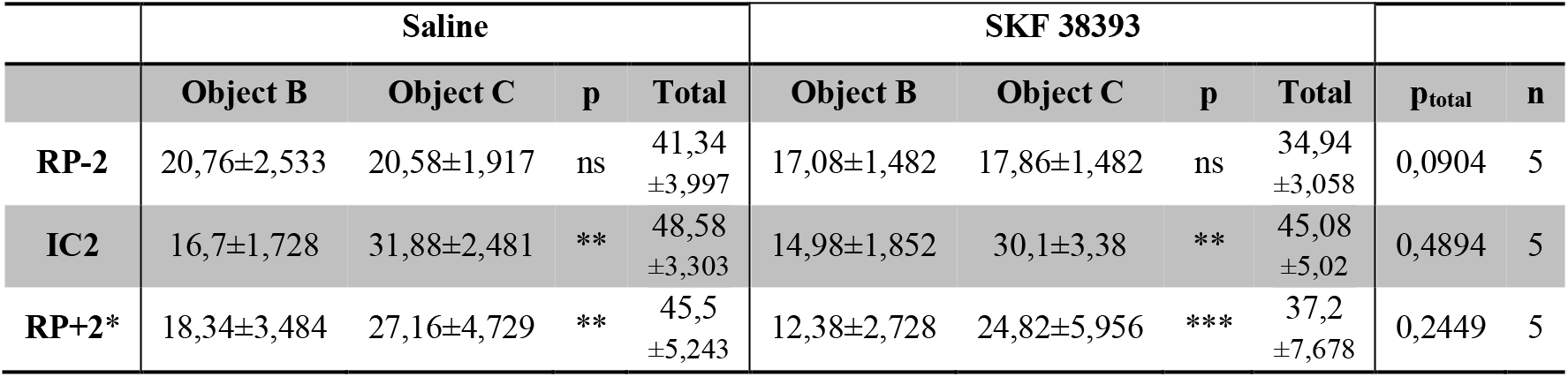
Exploration times during the final test phase for experiment depicted in Fig 3 A, (two practice sessions). **Absolute exploration time during the final test phase.** Total exploration times during the final test phase for the RP-2, IC2 and TC conditions. Values are expressed in seconds (mean ± S.E.M.). Paired student’s t test, comparing individual object exploration time between saline- and SKF-injected animals for the test phase; significance level is indicated as “p”. Paired student’s t test, comparing total exploration time between saline- and SKF-injected animals for the test phase (e.g SKF 38393 B+C mean vs saline B+C mean). Significance level is indicated as “p_total_”. *: p<0,05; **: p<0,01; ***: p<0,001; ****: p<0,0001.

|  | **Saline** |  |  |  | **SKF 38393** |  |  |  |  |  |
| --- | --- | --- | --- | --- | --- | --- | --- | --- | --- | --- |
|  | **Object B** | **Object C** | **p** | **Total** | **Object B** | **Object C** | **p** | **Total** | **p<sub>total</sub>** | **n** |
| **RP-2** | 20,76±2,533 | 20,58±1,917 | ns | 41,34<br>±3,997 | 17,08±1,482 | 17,86±1,482 | ns | 34,94<br>±3,058 | 0,0904 | 5 |
| **IC2** | 16,7±1,728 | 31,88±2,481 | ** | 48,58<br>±3,303 | 14,98±1,852 | 30,1±3,38 | ** | 45,08<br>±5,02 | 0,4894 | 5 |
| **RP+2\*** | 18,34±3,484 | 27,16±4,729 | ** | 45,5<br>±5,243 | 12,38±2,728 | 24,82±5,956 | *** | 37,2<br>±7,678 | 0,2449 | 5 |

**Figure 3.**
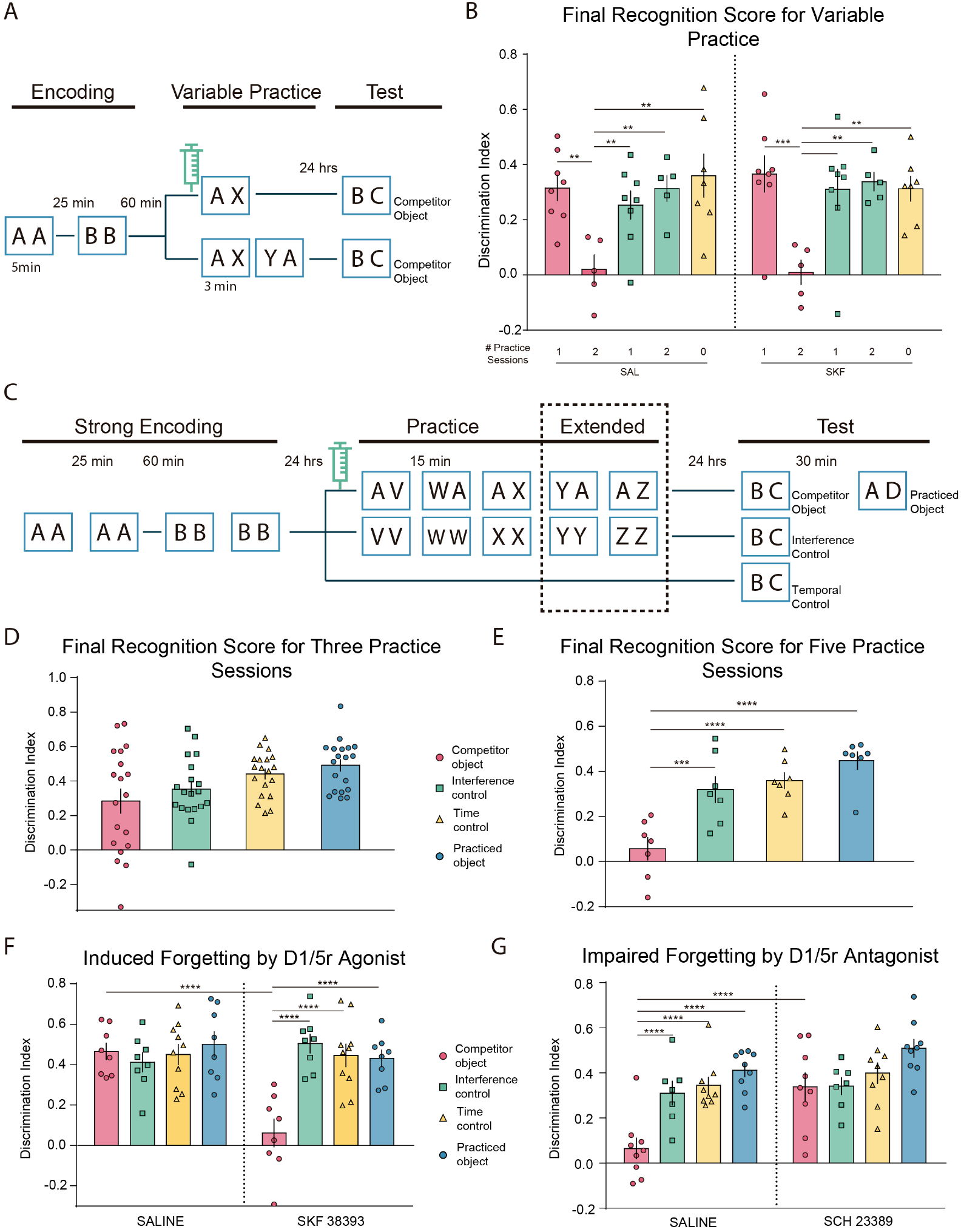
Bidirectional modulation of retrieval-induced forgetting. **(A)**. Schematic representation of the behavioral protocol. After the acquisition, the animals were divided into three conditions, RP, IC and TC. Both RP and IC were subdivided in two, a group that performed a practice phase with only one retrieval practice session (1) and another group that did two retrieval practice sessions (2). Only the RP group is schematized, the IC group performed the equivalent to the practice phase with two copies of identical objects (XX, or XX and then YY). The syringe indicates the infusion of SKF 38393 (SKF) or its vehicle (saline) 10 minutes before the practice phase. **(B)** Discrimination rates for the test phase. The animals did the task twice, once with the SKF and once with saline in a pseudorandomized way and for the same condition. Two-way ANOVA followed by a Bonferroni post hoc. **(C)** Schematic representation of the behavioral protocol. The protocol consisted of an acquisition phase with double training for each object (strong acquisition). After the acquisition, the animals were divided in three conditions, RP, CI and CT, the upper panels **(D and E)** correspond to two groups of animals that performed the protocol without infusion of any drug, the lower panels **(F and G)** correspond to other two groups of animals that were cannulated and infused with the D1R agonist and antagonist. The syringe indicates the infusion of the drug or its vehicle 10 minutes before the practice phase. (extended practice, one-way ANOVA for D and E, and two-way ANOVA for F and G; * p<0.05, ** p<0.01, ***p<0.001 and ****p<0.0001.

In this modified protocol, both IC and TC groups showed significant memory for the objects 48 hs after encoding (Figure 3D, Table 12). This longer delay succeeded in reducing retrieval-induced forgetting in the Veh group: Memory for the competitor object B in the RP group injected with Veh was not significantly different to that of the IC or TC groups after three practice sessions (Figure 3D, Table 12; one-way ANOVA: Condition: p= 0,2062, F(1,983, 35,7)= 4,055. Animals: p= 0,3591, F(18, 54)= 1,121; multiple comparisons). Critically, however, injection of the D1R agonist SKF into mPFC 15 min before the beginning of the retrieval practice session produced a robust memory impairment for the competitor object compared with the control groups (Figure 3F, Table 14; two-way ANOVA: Interaction: p< 0,0001, F(2, 23)= 25,50, Drug: p= 0,0016, F(1, 23)= 12,85, Condition: p= 0,013, F(23, 23)= 3,413. Bonferroni *post hoc* multiple comparisons). Thus, SKF amplified the capacity of the mPFC to hinder competing memories, enabling retrieval-induced forgetting even after 48 h.

**Table 12:** Exploration times and discrimination indexes during final test for experiment depicted in Fig 3 C, (normal practice phase). **Absolute exploration time during the final test.** Total exploration scores during the test phase for the RP-, IC, TC and RP+ conditions. Values are expressed in seconds (mean ± S.E). Within-subject experiment. Paired student’s t test, comparing individual object exploration time for the test phase; significance level is indicated as “p”. *: p<0,05; **: p<0,01; ***: p<0,001; ****: p<0,0001.

|  | Object B | Object C | p | DI | Total | n |
| --- | --- | --- | --- | --- | --- | --- |
| RP- | 19,68 ±2,085 | 37,78 ±3,624 | ** | 0,3027<br>±0,0631 | 57,46<br>±4,223 | 19 |
| IC | 16,87 ±1,42 | 35,98 ±2,532 | ** | 0,3526<br>±0,0421 | 52,85<br>±3,22 | 19 |
| TC | 14,81 ±1,122 | 37,69 ±2,119 | *** | 0,4409<br>±0,0295 | 52,5<br>±2,883 | 19 |
| RP+ | 13,08 ±1,269 | 37,03 ±2,565 | *** | 0,4915<br>±0,0334 | 50,11<br>±3,477 | 19 |

**Table 13:** Retrieval practice exploration times and discrimination indexes for experiment depicted in Fig 3 C, (normal practice phase with SKF 38393 infusion into the mPFC). **Absolute exploration time during the retrieval practice phase.** Total exploration times during the retrieval practice phase and DI for the RP and IC groups when animals were infused with SKF 38393 or saline in the mPFC. Values are expressed in seconds (mean ± S.E.M.). Paired Student’s t test, comparing total exploration time between saline- and SKF-injected animals for each retrieval practice session (e.g. Saline A+X mean vs SKF A+X mean for RP group and saline X_1_+X_2_ mean vs SKF X_1_+X_2_ mean, for IC group). Significance level is indicated as “p_total_”. SKF 38393 injection did not affect total exploration times during the practice phase compared to saline injection.

| RP | Saline |  |  | SKF 38393 |  |  | p total | n |
| --- | --- | --- | --- | --- | --- | --- | --- | --- |
|  | A | X/Y/Z | DI | A | X/Y/Z | DI |  |  |
| S 1 | 16,41<br>±2,953 | 24,76<br>±2,442 | 0,2185<br>±0,07098 | 15,8<br>±1,343 | 28,01<br>±3,89 | 0,2304<br>±0,07366 | 0,7408 | 8 |
| S 2 | 8,9<br>±0,8984 | 18,41<br>±4,329 | 0,3547<br>±0,08552 | 13,71<br>±2,878 | 18,18<br>±2,545 | 0,1709<br>±0,08577 | 0,5012 | 8 |
| S 3 | 10,86<br>±1,593 | 23,23<br>±5,493 | 0,4019<br>±0,0635 | 7,45<br>±1,048 | 23,39<br>±3,505 | 0,481<br>±0,07068 | 0,6835 | 8 |
| IC | X/Y/Z | X/Y/Z | DI | X/Y/Z | X/Y/Z | DI | p total | n |
| S 1 | 18,72<br>±2,782 | 19,87<br>±2,455 | 0,05167<br>±0,05501 | 16,38<br>±2,224 | 16,98<br>±1,586 | 0,03443<br>±0,04596 | 0,4248 | 8 |
| S 2 | 15,22<br>±2,471 | 16,68<br>±2,348 | 0,06551<br>±0,03051 | 15,39<br>±2,225 | 17,32<br>±1,946 | 0,07469<br>±0,05598 | 0,8892 | 8 |
| S 3 | 17,99<br>±4,772 | 18,52<br>±4,838 | -0,0059<br>±0,04875 | 15,31<br>±3,172 | 14,74<br>±2,877 | -0,03109<br>±0,03578 | 0,5892 | 8 |

**Table 14:** Exploration times during the final test phase for experiment depicted in Fig 3 C, (normal practice phase with SKF 38393 infusion into the mPFC). **Absolute exploration time during the final test phase.** Total exploration times during the final test phase for the RP-, IC,TC and RP+ conditions. Values are expressed in seconds (mean ± S.E.M.). Paired student’s t test, comparing individual object exploration time between saline- and SKF-injected animals for the test phase; significance level is indicated as “p”. Paired student’s t test, comparing total exploration time between saline- and SKF-injected animals for the test phase (e.g SKF 38393 B+C mean vs Saline B+C mean). Significance level is indicated as “p_total_”. *: p<0,05; **: p<0,01; ***: p<0,001; ****: p<0,0001.

|  | Saline |  |  |  | SKF 38393 |  |  |  |  |  |
| --- | --- | --- | --- | --- | --- | --- | --- | --- | --- | --- |
|  | Object B | Object C | p | Total | Object B | Object C | p | Total | p<sub>total</sub> | n |
| RP- | 11,84<br>±0,8878 | 34,04<br>±3,992 | *** | 45,88<br>±4,315 | 18,96<br>±2,318 | 22,95<br>±3,857 | 0,1814 | 41,91<br>±5,769 | 0,4718 | 8 |
| IC | 13,34<br>±1,586 | 34,02<br>±4,64 | *** | 47,37<br>±5,994 | 10,7<br>±1,367 | 34,03<br>±5,417 | ** | 44,73<br>±6,355 | 0,6967 | 8 |
| TC | 11,66<br>±1,659 | 33,6<br>±5,04 | *** | 45,26<br>±6,114 | 11,43<br>±1,447 | 35,93<br>±7,287 | ** | 47,37<br>±8,168 | 0,8122 | 9 |
| RP+* | 12,93<br>±1,47 | 33,58<br>±3,953 | ** | 48,01<br>±3,902 | 11,27<br>±1,054 | 36,74<br>±4,233 | *** | 46,51<br>±4,867 | 0,7543 | 8 |

To confirm that retrieval-induced forgetting could also occur in this longer protocol, we added two extra retrieval practice trials to the practice phase (see scheme in Figure 3C, ‘extended practice'). In Veh-injected rats, five retrieval practice trials induced significant memory impairment for the competitor object even at the 48 h post-encoding delay compared with matched IC and TC control groups (Figure 3E, Table 15, one-way ANOVA: Condition: p= 0,0002, F(1,984, 13,89)= 17,47. Animals: p= 0,0923, F(7, 14)= 2,257. Multiple comparisons). Injection of the D1R antagonist SCH into mPFC 15 min before the first of the 5 practice trials completely prevented forgetting of the competitor object, as performance was indistinguishable from the IC and TC groups (Figure 3G, Table 17; two-way ANOVA: Interaction: p< 0,0382, F(3, 30)= 3,18, Drug: p= 0,0009, F(1, 30)= 13,48, Condition: p< 0,0001, F(3, 30)= 11,51. Bonferroni *post hoc* multiple comparisons). Taken together, these show a bidirectional modulation of retrieval induced forgetting by manipulation of dopaminergic signaling through D1R in the mPFC.

**Table 15:** Exploration times and discrimination indexes during final test for experiment depicted in Fig 3 C, (extended practice). **Absolute exploration time during the final test phase.** Total exploration scores during the test phase for the RP-, IC, TC and RP+ conditions. Values are expressed in seconds (mean ± S.E). Within-subject experiment. Paired student’s t test, comparing individual object exploration time for the test phase; significance level is indicated as “p”. *: p<0,05; **: p<0,01; ***: p<0,001; ****: p<0,0001.

|  | Object B | Object C | p | DI | Total | n |
| --- | --- | --- | --- | --- | --- | --- |
| RP | 21,8 ±1,792 | 24,39 ±2,024 | 0,3486 | 0,0568<br>±0,04986 | 46,19 ±2,855 | 7 |
| IC | 13,17 ±1,639 | 26,3 ±2,899 | ** | 0,4482<br>±0,03919 | 43,34 ±3,869 | 7 |
| TC | 13,97 ±1,623 | 29,37 ±2,521 | *** | 0,3197<br>±0,05855 | 59,29 ±4,732 | 7 |
| RP+ | 16,03 ±1,015 | 43,26 ±4,029 | *** | 0,3588<br>±0,0357 | 39,47 ±3,86 | 7 |

**Table 16:** Retrieval practice exploration times for experiment depicted in Fig 3 C, (extended practice with SCH 23389 infusion into the mPFC). **Absolute exploration time and discrimination indexes during the retrieval practice phase for experiment 10.** Total exploration times during the retrieval practice phase and DI for the RP and IC groups when animals were infused with SCH 23389 or saline in the mPFC. Values are expressed in seconds (mean ± S.E.M.). Paired Student’s t test, comparing total exploration time between saline- and SCH-injected animals for each retrieval practice session (e.g. Saline A+X mean vs SCH A+X mean for RP group and saline X_1_+X_2_ mean vs SCH X_1_+X_2_ mean, for IC group). Significance level is indicated as “p_total_”. SCH 23389 injection did not affect total exploration times during the practice phase compared to saline injection.

|  | Saline |  |  | SCH 23389 |  |  |  |  |
| --- | --- | --- | --- | --- | --- | --- | --- | --- |
| RP | A | X/Y/Z | DI | A | X/Y/Z | DI | p total | n |
| S1 | 15,39<br>±1,642 | 26,68<br>±2,51 | 0,2656<br>±0,0620 | 12,14<br>±1,581 | 26,82<br>±2,434 | 0,2954<br>±0,0545 | 0,1044 | 9 |
| S2 | 14,36<br>±1,574 | 29,18<br>±3,142 | 0,3358<br>±0,0407 | 10,66<br>±1,342 | 24,07<br>±2,657 | 0,3719<br>±0,0412 | 0,1223 | 9 |
| S3 | 11,89<br>±1,724 | 23,24<br>±2,039 | 0,2856<br>±0,0743 | 8,633<br>±1,185 | 23,16<br>±2,458 | 0,2687<br>±0,0528 | 0,3970 | 9 |
| S4 | 11,14<br>±1,919 | 24,46<br>±4,337 | 0,3136<br>±0,0925 | 9,078<br>±1,723 | 21,81<br>±2,336 | 0,3204<br>±0,0694 | 0,3082 | 9 |
| S5 | 10,02<br>±1,407 | 25,16<br>±3,487 | 0,4126<br>±0,0432 | 7,956<br>±2,256 | 16,68<br>±2,415 | 0,4505<br>±0,0884 | 0,2150 | 9 |
| IC | X/Y/Z | X/Y/Z | DI | X/Y/Z | X/Y/Z | DI | p total | n |
| S1 | 19,71<br>±3,053 | 21,45<br>±2,043 | 0,0744<br>±0,0660 | 16,24<br>±2,909 | 18,89<br>±2,056 | 0,1140<br>±0,0465 | 0,5706 | 7 |
| S2 | 19,41<br>±1,888 | 19,06<br>±1,395 | 0,0226<br>±0,0182 | 17,09<br>±1,635 | 18,15<br>±2,492 | 0,0375<br>±0,0462 | 0,7549 | 7 |
| S3 | 15,79<br>±1,747 | 15,03<br>±2,154 | -0,0335<br>±0,0245 | 9,575<br>±2,328 | 13,01<br>±1,654 | 0,0589<br>±0,1273 | 0,2798 | 7 |
| S4 | 14,39<br>±2,770 | 15,14<br>±3,101 | -0,0256<br>±0,0438 | 9,388<br>±1,439 | 10,69<br>±3,038 | -0,0152<br>±0,0864 | 0,1411 | 7 |
| S5 | 12,89<br>±2,834 | 11,73<br>±2,834 | -0,0022<br>±0,0658 | 10,81<br>±2,710 | 13,21<br>±2,498 | 0,1081<br>±0,0962 | 0,9886 | 7 |

**Table 17:** Exploration times during the final test phase for experiment depicted in Fig 3 C, (extended practice with SCH 23389 infusion into the mPFC). **Absolute exploration time during the final test phase.** Total exploration times during the final test phase for the RP-, IC, TC and RP+ conditions. Values are expressed in seconds (mean ± S.E.M.). Paired student’s t test, comparing individual object exploration time between saline- and SCH-injected animals for the test phase; significance level is indicated as “p”. Paired student’s t test, comparing total exploration time between saline- and SCH-injected animals for the test phase (e.g SCH 23389 B+C mean vs Saline B+C mean). Significance level is indicated as “p_total_”.*: p<0,05; **: p<0,01; ***: p<0,001; ****: p<0,0001.

|  | Saline |  |  |  | SCH 23389 |  |  |  |  |  |
| --- | --- | --- | --- | --- | --- | --- | --- | --- | --- | --- |
|  | Object B | Object C | p | Total | Object B | Object C | p | Total | p<sub>total</sub> | n |
| RP- | 18,72 ±1,985 | 21,38<br>±1,850 | 0,0661 | 40,1<br>±3,628 | 14,36<br>±1,129 | 30,7 ±3,502 | ** | 45,06<br>±3,877 | 0,3298 | 9 |
| IC | 14,47 ±0,918 | 29,19<br>±4,578 | * | 43,66<br>±5,031 | 11,57<br>±1,305 | 23,9 ±2,58 | *** | 35,47<br>±3,739 | 0,2500 | 7 |
| TC | 13,38 ±1,182 | 28,41<br>±2,869 | *** | 41,79<br>±3,645 | 12,63<br>±1,072 | 30,22<br>±2,819 | *** | 42,86<br>±3,342 | 0,8368 | 9 |
| RP+ | 13,00 ±1,161 | 31,07<br>±2,027 | **** | 44,07<br>±2,872 | 10,8<br>±1,575 | 31,58<br>±4,059 | *** | 42,38<br>±5,09 | 0,7631 | 9 |

## Discussion

Memory is essential to adaptive behavior. It enables organisms to draw on past experience to improve choices and actions. Because of their relational nature and richness, episodic memories are flexible in a way that past events can be retrieved as needed to guide future behavior (Eichenbaum et al., 2001). Experience has been shown to modify behavior in several ways by restructuring access to memories or directly modifying the memory traces (Quirk and Mueller, 2008; Lee, 2009; Medina, 2018). The neurotransmitter dopamine plays an important function in the ability to change a learned rule and select appropriate behaviors (Seamans and Yang, 2004) by biasing action selection and even modifying neural plasticity in regions of memory storage (Lisman and Grace, 2005; Neugebauer et al., 2009). Thus, dopamine is considered an important player for adaptive behavior. In this work, we expand the functions of dopamine to a mechanism of adaptive and selective forgetting of competing memories. Although the role of dopamine has been mainly studied in motivation of goal-directed behaviors, we argue that dopamine-dependent mechanisms are related to general adaptive memory processing even in the absence of any type of explicit reward. We propose that retrieval-induced forgetting of a neutral competing object memory operates under similar mechanisms than that of rule switching and selection in the mPFC of rodents and that this modulation of control processes help to adapt memory content to the behavioral demands of the organism. Retrieval-induced forgetting in rats remarkably resembles this process in humans. Critically, the mPFC in rats is essential to forget competing object memories, paralleling results observed for the lateral prefrontal cortex in humans. These results point to the key role of inhibitory control processes as an essential part of retrieval-induced forgetting. In this study we provide strong causal evidence in favor of a dopamine-dependent mechanism of inhibitory control for retrieval-induced forgetting. Blockade of D1R in the mPFC of rats during the practice phase completely prevented retrieval-induced forgetting of an object competing memory. This manipulation did not have any effect when it preceded the encoding of novel interfering materials (Interference control) or when it preceded rest in the rats’ home cage, indicating that it affected processes specifically associated with retrieval practice. The function of D1R in the prefrontal cortex has been extensively investigated. Many studies have found that D1R blockade in non-human primates disrupts task performance and spatial working memory activity in the dorsolateral prefrontal cortex (DLPFC) (Sawaguchi and Goldman-Rakic, 1991, 1994; Williams and Goldman-Rakic, 1995). Importantly, D1R blockade also disrupts prefrontal cognitive rule-related selectivity (Ott et al., 2014). In this work, we found that the same dose and place of infusion of the D1R antagonist that prevented retrieval-induced forgetting also impaired performance in a set shifting task in which rats are required to inhibit a prepotent response associated to a learned rule. The parallel impact of a D1R antagonist on the need to inhibit prepotent actions and memories is consistent with human studies indicating that retrieval-induced forgetting is triggered by inhibitory control processes shared with action stopping (Schilling et al., 2014; Anderson and Hulbert, 2021). Also, it provides new evidence in favor of a general function of dopamine in cognitive processes related to flexible and adaptive behavior.

We provided causal evidence that the critical source of dopamine for retrieval-induced forgetting in the mPFC is the VTA, because silencing this structure impaired retrieval-induced forgetting. This effect was reversed by concomitant activation of D1R in mPFC during the practice phase indicating that, in the absence of dopamine release from VTA, activation of D1R in the mPFC is sufficient for retrieval-induced forgetting. Critically, dopaminergic modulation of retrieval-induced forgetting is bidirectional. Activation of D1R in the mPFC just before the retrieval practice phase caused retrieval-induced forgetting in a protocol that does not reliably induce it without D1R activation. No anxiety, movement or perception changes were observed after any of the infusions, as rats did not significantly modify their exploratory behavior after infusion of any of the drugs.

The strong link between dopamine availability in the brain and cognitive abilities has been long known. Interestingly, many of the studies point at a function of dopamine in adaptive behavior in humans. For example, administration of L-DOPA to Parkinson’s disease patients has been reported to improve the ability to alter behavior according to changes in dimensional relevance of stimuli in a task that resembles the set-shifting paradigm used in our study (Cools et al., 2001). Interestingly, the impairments in this form of higher-level attentional control have been associated with lesions of the monkey lateral PFC (Dias et al., 1996) and significant activation of the DLPFC in humans (Rogers et al., 2000; Nagahama et al., 2001). In addition, the enzyme COMT, which degrades catecholamines, appears to play the pivotal role in the modulation of fronto-striatal networks. Many studies report an association between the COMT Val158Met polymorphism and cognitive function. The *COMT* gene presents an evolutionary recent functional single nucleotide polymorphism (SNP) (Val158Met). The Met allele produces an enzyme that has only a quarter the activity of the Val-containing polypeptide (Egan et al., 2001). These studies suggest that the low-activity Met allele allows for better performance on cognitive tasks that have a working memory component and the high-activity Val allele was associated with poorer performance on the Wisconsin Card Sorting Test (WCST), a putative measure of ‘executive’ function (reviewed in (Savitz et al., 2006)). Interestingly, in humans, retrieval-induced forgetting increased linearly with Met allele load, suggesting a positive relationship between cortical dopamine availability and inhibitory control over memory (Wimber et al., 2011). Mirroring the linear effect of genotype on behavior, functional imaging data revealed that the beneficial effects of memory suppression, as assessed by a decrease in prefrontal brain activity across retrieval practice blocks, a sign of efficient suppression of competing memories (Kuhl et al., 2007; Bekinschtein et al., 2018; Anderson and Hulbert, 2021), also increased with Met allele load. In agreement with these results, the present study supports a general contribution of dopamine in the mPFC in both control processes and, in particular, establishes causality between dopamine availability and retrieval-induced forgetting. Greater dopamine availability in this structure may lead to greater activation of D1R receptors improving suppression of competing memories. Efficient suppression of competing information should lead to better memory performance and more adaptive behavior.

What are the mechanisms by which dopamine participates in retrieval-induced forgetting? Activation of D1R in mPFC could initiate active circuit-level inhibition over competing memory traces in the medial temporal lobe. Given that top-down connections from the mPFC to the medial temporal lobe are mainly excitatory (Hoover and Vertes, 2007; Vertes et al., 2007) projections from the mPFC would not directly enact inhibition over the competing memory trace. A possible mechanism could involve excitatory projections from the prefrontal cortex that directly excite local inhibitory neurons in the site to be influenced, which then inhibit a distracting stimulus, or unwanted representation or process (Chamberland and Topolnik, 2012). The rodent infralimbic cortex originates modest projections to the entorhinal and ectorhinal (analogous to perirhinal area 36 in macaque monkeys) cortices, while pathways originating from prelimbic cortex specifically target the entorhinal cortex (Vertes, 2004). We hypothesize that the mPFC-MTL pathway can invoke memory retrieval inhibition of competing memories. If the competing memory trace was stored in the hippocampus, the activation of the mPFC could induce inhibition of the competing traces in the hippocampus via nucleus reuniens (RE) (Vertes, 2006), since there are no direct connections from the mPFC to this region. This ‘RE hypothesis’ would implicate that excitatory projections from the mPFC in modulating the activity of inhibitory neurons in the hippocampus where the competing memory trace would be suppressed (Anderson et al., 2016). Although this remains speculative, recent evidence suggests that projections from mPFC to RE play a role in modulating excitability of hippocampal neurons, thereby controlling the specificity with which memories are encoded. Alterations to RE-hippocampal interactions influence the tendency to overgeneralize fear memories to novel contexts in which fearful events did not happen (Xu and Südhof, 2013; Ito et al., 2015), a tendency that maybe relevant to contextually inappropriate recall of traumatic flash-back memories. We propose that dopamine modulates mPFC activity that, in turn, increases or decreases activation of neurons in the RE that project to the hippocampus and affects local inhibitory interneurons involved in retrieval-induced forgetting(Anderson et al., 2016).

Regardless of the circuit involved in retrieval-induced forgetting, we made the surprising discovery that dopaminergic modulation of retrieval-induced forgetting seems to be independent of any mechanisms of retrieval itself (i.e., D1R blockade in mPFC does not affect retrieval during the practice phase but impairs retrieval-induced forgetting). This suggests that dopamine modulates retrieval-induced forgetting by specifically acting on the future availability of the competing memory trace (i.e., at the test phase), without affecting the retrieval processes during the practice phase. Thus, we argue that retrieval control and retrieval-induced forgetting mechanisms are intrinsically distinct. During retrieval practice, activity in the mPFC would be required for later impairment in the retrieval of the competing memory, but not for the mechanism of retrieval itself. Lesions to the mPFC in rats do not normally impair object recognition when the task relies on the identity of the object (Warburton and Brown, 2015). However, what we found is that even if the mPFC is not implicated in object memory retrieval, it does not mean that the structure does not participate in memory at all. The high-level functioning of the mPFC would allow for adaptive memory processing relying on previous experience. In particular, D1R would be essential for this high-level function.

In agreement with an adaptive and evolutionary conserved role in memory and behavior, dopamine has been recently implicated in forgetting mechanisms in both invertebrates (Berry et al., 2012) and vertebrates (Wimber et al., 2011; Castillo Diaz et al., 2019). Modulation of a small subset of dopaminergic neurons in *Drosophila* regulates the rate of forgetting of aversive and rewarding experiences. In particular, forgetting appears to depend on signaling through a specific type of receptor in the mushroom bodies of the fly brain (Berry et al., 2012). On the other hand, inhibition of D1R in the VTA during training of a conditioned place preference task in rats, increases memory duration, while activation of these receptors produces forgetting of already consolidated memories (Castillo Diaz et al., 2019). In the absence of any type of retrieval practice, blockade of mPFC D1R did not produce forgetting of the conditioned place preference memory. Although this study did not evaluate the function of D1R in retrieval-induced forgetting it does contribute to an increasing accumulation of evidence for the involvement of the dopaminergic system in the different mechanisms of forgetting linked to adaptive behavior.

According to our results, dopamine acting on D1R in the mPFC modulates control processes required for adaptive forgetting in the mammalian brain. Thus, across species, dopaminergic transmission may be essential to suppress competing memories by sculpting the mnestic and behavioral repertoire of an organism according the demands of the environment.

## Materials and Methods

### Subjects

245 male adult Wistar rats (weight 180-250 gr and between 8 and 12 weeks old) were housed socially up to five per cage and kept with water and food *ad libitum* under a 12 h light/dark cycle (lights at 7 A.M.) at a constant temperature of 23 °C. Separate groups of animals were used for the different experiments. Experiments took place during the light phase of the cycle. The experimental protocol for this study followed guidelines of the National Institute of Health Guide for the Care and Use of Laboratory Animals and government regulations (SENASAARS617.2002) and the law 14346 on animal protection. It was approved by the institutional Committee for care and use of laboratory animals of Universidad Favaloro (UF DCT0204-16) and the Institutional Animal Care and Use Committee of the School of Medicine, University of Buenos Aires (ASP # 49527/15). The number of animals used is stated for each experiment (see below). All efforts were made to minimize the number of animals used and their suffering.

### Apparatus

Different arena contexts were used during the experiments.

Our experiments are, mostly, *within-subjects drug designs*, or *within-subjects behavioral designs*. All animals were exposed to at least 4 contexts during the experiment in which they participated. Animals in the within-subjects behavioral designs were exposed to a total of 6 contexts. Except for contexts 5, 7 and 8, that were used exclusively to habituate animals to the objects presented as contextually novel during the practice phase, all other contexts were assigned pseudo-randomly to each phase of the experiment. All animals that take a variation of the retrieval practice paradigm went through a shaping phase (see explanation below) and then started the experiment.

Arena 1 was a 50 cm wide × 50 cm long × 39 cm high arena with black plywood walls and floor, divided into 9 squares by white lines.
Arena 2 was a 60 cm wide × 40 cm long × 50 cm high acrylic box. The floor was white as well as two of its walls, which had different visual cues, geometric forms or strips made with self-adhesive paper tape of different colors. The frontal wall was transparent and the back wall was hatched.
Arena 3 was a 50 cm diameter × 50 cm high round arena with brown acrylic walls and black plywood floor, divided into 9 squares by white lines.
Arena 4 was a 50 cm wide × 50 cm long × 40 cm high box constructed with white Plexiglas. The floor was made of white Plexiglas as well. Each wall had different visual cues, geometric forms or strips made with self-adhesive paper tape of different colors.
Arena 5 was a 40 cm diameter × 50 cm high round arena with brown acrylic walls and sky-blue floor.
Arena 6 was a bow-tie-shaped maze made of opaque white Plexiglas. The maze was 94 cm long, 50 cm wide, and 50 cm high. Each end of the apparatus was triangular, the apexes of which were joined by a narrow corridor (14 cm wide).
Arena 7 was a Y-shape apparatus constructed from Plexiglas. All walls were 40 cm high, and each arm was 27 cm in length and 10 cm wide.
Arena 8 was a equilateral triangular arena of 40 cm side × 40 cm high made of white semi-rigid PVC with white floor made of the same material.

### Objects

All experiments used numerous junk objects, each differing in shape, texture, size, and color. The height of the objects ranged from 8cm to 24 cm and they varied with respect to their visual and tactile qualities. All objects had duplicates so that identical objects could be used at the same time. All objects were affixed to the floor of the apparatus with an odorless reusable adhesive to prevent them for being displaced during each session. Specific objects were never repeated across different conditions for a given animal. All objects were cleaned with 50% alcohol wipes after each session.

### Memory Test for Retrieval-Induced Forgetting

#### Overview

We modified the spontaneous object recognition task to study retrieval-induced forgetting (RIF, for details see(Bekinschtein et al., 2018)). For each experiment, different cohorts of animals were used. The order in which they were exposed to each condition (or drug/vehicle) was pseudo-randomly assigned and the three conditions were conducted over a span of three weeks (or two weeks for the within-subject drug design). Once we finished evaluating the animal for one of the conditions (e.g. Retrieval Practice), we waited 3 days to start testing the following condition (e.g. Interference Control). For the drug design, animals waited at least 4 days.

#### The general retrieval practice paradigm

Our new retrieval practice paradigm generally involved three conditions: the Retrieval Practice (RP), Interference Control (IC) and Time Control (TC) conditions (Bekinschtein et al. 2019). Every condition followed the same basic sequence across three days: *Day 1*: habituation to the contexts, *Day 2*: Habituation to “distractor” objects to be used during the retrieval practice phase of the experiment, *Day 3*: the main memory task. During the main memory task, encoding and practice phases took place in a single session, and *Day 4*: test phases.

#### Habituation

We incorporated a shaping procedure that included four sessions of object exposure. During this shaping, rats were first habituated to two different contexts (10 minutes each, not described in *Apparatus*), three hours later rats were exposed to two pairs of novel objects in two contexts. The animals were exposed twice to each context (four sessions) with a delay of 20 minutes. In each session that lasted 5 min, the rats encountered the same two pairs of different objects in distinct locations. The objects were novel during the first exposure, but familiar during the next three. Each rat saw the four objects twice in both contexts. For each context, the location of the objects was different between the first and the second exposure. The shaping phase was conducted only once during the first week of the experiment independently of the condition assigned for that particular week. We added this procedure to familiarize rats with the possibility that the very same objects could be presented in different locations within a context or across contexts (Bekinschtein et al. 2019). All experiments started 72 hours after shaping.

On the first day of the experiment, animals were habituated to two arena contexts (e.g.: contexts 1 and 2) and allowed to explore each context for 10 minutes. On the second day, each animal was exposed to three pairs of identical novel objects (X, Y and Z) in context 2 in three consecutive (30 min apart) sessions, for 5 min each. The following day, the task was conducted in context 1.

#### Retrieval Practice (RP) condition

The sample phase consisted of two consecutive sessions separated by 30 minutes. In these sample sessions, the animal was allowed to freely explore for 5 min two identical copies of two novel objects: e.g., object A (session 1) and object B (session 2). The practice phase took place thirty minutes after the last sample session. This phase consisted of three 3-min sessions with an intersession interval of 15 min. In each session the animal was exposed to a copy of one of the two encoded objects (e.g., Object A) presented during the sample phase--accompanied by one copy of objects X, Y or Z respectively across the three trials (e.g., A & X; then A& Y; then A& Z across the three sessions). We pseudo-randomly assigned which object was presented during the retrieval practice phase from the two objects that were sampled in the sampling phase (either A or B), so the practiced object could either be the first or the second one that was encoded in the sampling phase. Moreover, the location (right or left) in which the studied object appeared during retrieval practice was randomly assigned for each trial. The test phase was conducted 24 hours after the last practice session. The animal was exposed for 3 min to a copy of a non-practiced competitor object presented only during the sample phase (e.g., Object B) and one completely novel object never before seen (object C). Twenty minutes later the animals were re-introduced to the context and exposed for 3 minutes to a copy of practiced object (object A) and one completely novel object (object D). These two test sessions are defined in the results section as “competing object” and “practiced object”, respectively. For both test sessions the location of the novel and familiar objects (right or left) was randomly assigned. The letters used in these descriptions and in our diagrams and meant to identify indicate the nature of the item--practiced object, competitor object, novel object or distractor. Repetitions of the same letter across conditions do not indicate that the same object was used across conditions: in fact, different objects were used for the different conditions – RP, IC or TC- of the task. Thus, object A used in the RP condition is different from object A used in the IC or TC conditions.

#### Interference Control (IC) condition

On the first day, the animals were habituated to two contexts (e.g., contexts 3 and 4) and allowed to explore them for 10 minutes each. On the second day, each animal was exposed to three novel objects (X, Y and Z) in three consecutive (30 min apart) sessions, in context 4 for 5 min each. On the third day, the main memory task was conducted in context 3. On this final day, during the sample phase each rat was allowed to freely explore for 5 min two identical copies of two novel objects (objects A and B) in two consecutive sessions separated by 30 minutes. The practice phase took place thirty minutes after the sample phase. During this phase, the animal was allowed to explore two copies of objects X, Y and Z in context 3 during three consecutive 3-min sessions with a delay of 20 min between each session. The test phase (30 min after the last practice session) consisted of a 3-min exposure to a copy of object B and one completely novel object (object C). Twenty minutes later the animals were re-introduced into the context and exposed for 3 min to a copy of object A and one completely novel object (object D). The time the animals spent exploring the objects in each trial was manually recorded using hand chronometers. The order in which the sample objects were tested was pseudo-randomly assigned and the position in which the sample objects appeared on the final test was randomly determined.

#### Time Control (TC) condition

On the first day, the animals were habituated to one context (e.g.: arena context 5), and allowed to explore it for 10 minutes. On the second day, the animals were transferred to the behavioral testing room but allowed to stay in their home cage for the duration of time that the animals assigned to the other two conditions were habituated to the novel objects. On the third day, the main memory task was conducted in context 5. The sample phase consisted of two consecutive sessions separated by 30 minutes. In these sessions, the animal was allowed to explore freely for 5 min two identical copies of two novel objects A (session 1) and B (session 2). Unlike in the RP and IC conditions, however, there were no practice trials; instead, the rats spent the same interval of time in their home cages in between the sample phase and the test. The test phase took place at the end of this two-hour interval; during this phase, the animal was exposed to a copy of object B and a completely novel object (object C) for 3 min. Twenty minutes later the animals were re-introduced to the context and exposed for 3 minutes to a copy of object A and one completely novel object (object D). The order in which the sample objects were tested was pseudo-randomly assigned and the position in which the sample objects appeared on the final test was randomly determined.

#### Quantification of behavior

The behavioral responses of the animals for all experiments were analyzed given the following criteria. We defined exploration of an object as the rat directing its nose to the object at a distance of <2 cm and/or touching it with its nose. Turning around or sitting on the object was not considered exploratory behavior. Encoding, practice and test phases were recorded using Samsung HMX-F80 cameras. The cameras were located on top of each arena allowing the visualization of the complete space. Offline analysis was done using the Stopwatch software (Center for Behavioral Neuroscience, Emory University) by a trained person. The test phase was analyzed by an experimenter who was blind to the conditions of the experiment.

Based on these criteria, we calculated a discrimination index (DI) for each trial of each session on each condition, as follows:

##### Practice trials

a discrimination index was calculated as the difference in time spent exploring the contextually novel and familiar objects divided by the total time spent exploring the objects (i.e. [(contextually novel – familiar)/total exploration time]).

##### Test trials

a discrimination index was calculated as the difference in time spent exploring the novel and familiar objects divided by the total time spent exploring the objects (i.e. [(novel – studied)/total exploration time]).

#### Criteria of exclusion

Animals that explored the objects for less than 10 sec during any of the phases would be excluded from the experiments. However, no rats had to be excluded from the study based on this criterion. Once the animals recovered from surgery the behavioral procedure started. Rats were run in groups of 8–10 per week and randomly assigned to each experimental group at the beginning of the experiment. So, all conditions within an experiment were run simultaneously.

### Specific design features of individual experiments

#### Surgery and Drug Infusions

Animals were habituated to experimental manipulation before surgery. Movements and positions required for future intracranial injection were performed. The day of surgery, animals were habituated at least one hour to the room where pharmacological anesthesia was injected. Rats were deeply anesthetized with ketamine (60 mg/kg) and xylazine (8 mg/kg) and put in a stereotaxic frame (Stoelting). The skull was exposed and adjusted to place bregma and lambda on the same horizontal plane. After small burr holes were drilled, a set of 22 g guide cannulas were implanted bilaterally into the mPFC (AP +3.20 mm/ LL ± 0.75 mm / DV −3.50 mm) and/or the VTA (AP −7.20 mm / LL ± 0.75 mm / DV −5.30 mm) (Paxinos & Watson, 1998). Cannulae were fixed to the skull with dental acrylic. A dummy cannula was inserted to each cannula to prevent clogging. At the end of surgery, animals were injected with a single dose of meloxicam (1 mg/kg) as an analgesic and gentamicin (0.6 mg/kg) as antibiotic.

Rats were allowed to recover for 7 days before starting any behavioral experiment. During these days, the animals were monitored by cageside observation of body posture and activity level and if any sign of discomfort (e.g. sickness, infection of the wound, loss of body weight) was displayed, special care was administered (e.g., extra dose of analgesic, subcutaneous injection of saline, antibiotic treatment). Before the behavioral experiment itself, a round of “shaping” was conducted, which include habituation to the type of task and to the experimental manipulation. If at the moment of the intracranial injection the animal showed clear signs of pain, it was not included in the trial. On the experimental day, the dummy cannulas were removed before the injection and an injection cannula extending 1 mm below the guide cannula was inserted. The injection cannula was connected to a 10 μl Hamilton syringe. Cannulated rats received bilateral 0.5 μl infusions the corresponding drug/vehicle. Muscimol (0.1 mg/ml in saline, Sigma #2763-96-4) infusions into the VTA occurred 15 minutes before the retrieval practice phase (or at the corresponding points in TC conditions). SCH 23389 (3 mg/ml in saline, Tocris #0925/10) and SKF 38393 (8.41 mg/ml in saline, Tocris #0922/100) occurred 10 minutes before the retrieval practice phase (or at the corresponding points in TC conditions). We conducted the final test 24 h later.

### Cannula placement

To check cannula placement, 24 h after the end of the behavioral experiments, animals were infused with 1 μl of methylene blue through the dummy cannulae and 15 min later deeply anesthetized and sacrificed. Histological localization of the infusion sites was established using magnifying glasses. 5 animals were excluded because of cannulae misplacement confirmed with the infusion of Green Beads (Lumafluor Inc.).

### Experimental design and Statistical analysis

Statistical analyses were performed using GraphPad 6.01. Behavioral data were analyzed using two-tail unpaired Student’s t test when two groups were compared. For comparisons between two repeated-measured groups, two-tail paired Student’s t test was used. One- or Two-way ANOVA followed by Bonferroni post-test, as indicated in the figure legends, was used when three or more groups were involved. In all cases, p values were considered to be statistically significant when p<0.05. Unless otherwise stated, p-values indicated in the figure footnotes refer to multiple comparisons. All data is presented as the mean ± s.e.m. The general retrieval practice paradigm is designed to study the behavioral effect of the experimental conditions in a within-subject approach. Thus, all animals perform all experimental conditions. The general retrieval practice paradigm experiments where we did not use drug infusions met this condition (Figures 3D and 3E). In the general retrieval practice paradigm experiments where we used drug infusions the “drug” variable was analyzed in a within-subject approach and the “condition” variable in a between-subjects approach. Experiments that met this condition are Figures 1C, 2C, 3B, 3F and 3G. For data details see *Supplementary tables*.

### Set Shifting task

#### Apparatus

The cross-maze was a four-arm maze made of 1-cm-thick black plexiglass (see Figure 1F). The maze was placed on the floor. Each arm was 52 cm long and 9 cm wide; the height of the arm wall was 40 cm. Each arm contained a food well (3 cm diameter, 2.5 cm high) that was 3.2 cm from the end wall.

#### Habituation Procedure

The habituation procedure was similar to that described in (Ragozzino, 2002). Rats were allowed 7–10 d to recover from surgery before the habituation procedure commenced. Rats were food restricted to 85% of their original *ad libitum* weight. During food restriction rats were handled for 10 min per day. On the first day of habituation, 3 pieces of Fruit Loops cereal (Kelloggs) were placed in each arm, with 2 pieces in the food well. A rat was placed in the maze and allowed to freely navigate and consume cereal pieces for 15 min. If a rat consumed all 12 cereal pieces prior to 15 min, then the rat was placed in a holding cage, the maze was rebaited, and the rat was placed back in the maze; this process was repeated a total of 3 times (if a rat did not consume all 12 cereal pieces prior to 15 min, then the habituation day 1 was repeated the next day until the rat reach criterion). On the second habituation day, the procedure was similar except that after a rat consumed 2 cereal pieces per arm, the rat was picked up and placed in a different arm. This acclimated the rat to being handled in the maze after consuming cereal. On subsequent habituation sessions, the procedure was the same as day 2, except that there were only 2 half pieces of cereal put in each food well. Each time a rat consumed all the cereal pieces after being placed in the maze was considered one trial. This procedure continued until a rat consumed cereal from all food wells for four trials or more in a 15-min session. On the last day of habituation, the turn bias for a rat was determined. The maze was arranged such that a white Plexiglas block (9 × 40 × 1 cm) was placed at the center entrance of one of the arms so that it prevented entry into that arm, giving the maze a T-shape. A rat was started from the stem arm and allowed to turn left or right to obtain a half piece of cereal. In one of the choice arms a white blue piece of posterboard (8 × 48 × 0.3 cm) was placed on the floor (see Fig. 1F). After a rat made a turn and consumed a cereal piece, the rat was picked up, placed in the stem arm, and allowed to make a choice. If the rat chose the same arm as in the initial choice, it was returned to the stem arm until it chose the other arm and consumed the cereal piece. After choosing both arms, the rat was returned to the holding cage, the block and visual cue were moved to different arms, and a new trial was begun. Thus, a trial for the turn-bias procedure consisted of entering both choice arms and consuming both cereal pieces. This procedure continued for seven trials. The turn that a rat made first during the initial choice of a trial was recorded and counted toward its turn bias. Whatever direction (right or left) a rat turned, four or more times during these seven trials was considered its turn bias. During response-discrimination testing, a rat was required to turn in the opposite direction of its turn bias. Behavioral testing was started the next day.

#### Response–Visual-Cue Testing Procedure

The testing procedure was similar to that described in Ragozzino et al. (2002) (Ragozzino, 2002) except that all testing was carried across two consecutive sessions. For each discrimination, three start arms were used. In this experiment, each rat was started on the response version. A rat was started from the arms designated west, south, and east (W, S, and E, respectively) leaving the north arm unused as starting arm. The visual cue was placed pseudo-randomly in one of the choice arms such that for every consecutive set of 12 trials it occurred an equal number of times in each choice arm. During the acquisition session, a rat had to turn in the opposite direction of its turn bias to receive a half piece of Froot Loops cereal. Figure 1F (top) illustrates an example of the correct navigation patterns for a rat that was required to always make a turn to the right. Between trials a rat was placed back in the holding cage, which sat on a shelf next to the maze. The intertrial interval was less than 20 sec. To minimize the use of intra-maze cues from the apparatus, every 6 trials the maze was turned 90° clockwise relative to the experimenter. A rat reached criterion when it made 10 correct choices consecutively. There was no limit on the number of trials a rat was prearranged to reach this criterion. Once a rat made 10 correct choices consecutively, a probe trial was given. The probe trial consisted of starting the rat from the fourth arm (north, N) that was not used during testing. If a rat correctly turned the same direction as on testing, then the response procedure was completed. If a rat made an incorrect turn, then response testing was continued until a rat made an additional 5 correct choices consecutively, at which time another probe trial was administered. This procedure was continued until a rat made a correct choice on the probe trial. The following measures were taken for each rat: (1) Acquisition criterion, defined as the total number of test trials to complete 10 consecutive correct choices in a session; (2) Trials to criterion, defined as the total number of test trials completed before a correct choice on the probe trial was made; and (3) Probe trials, defined as the total number of probe trials to get one correct. The day after reaching criterion on the response version, rats were switched to the visual-cue version. In the visual-cue version a similar procedure was used as in the response version. However, in this test the rat always had to enter the arm with the visual cue. The visual cue was pseudo-randomly varied in the left and right arms such that it occurred in each arm an equal amount for every consecutive set of 12 trials. Figure 1F (*bottom*) shows an example of a rat that learned to always enter the visual-cue arm. A rat reached criterion when it made 12 correct choices consecutively. There was no limit on the number of trials a rat was allotted to reach this criterion. Once a rat made 12 correct choices consecutively, a probe trial was given. If a rat correctly turned following the visual cue, then the response procedure was completed. If a rat made an incorrect turn, then visual testing was continued until a rat made an additional 6 correct choices consecutively, at which time another probe trial was administered. Additional measures were analyzed on the switch to determine whether treatments altered perseveration. Perseveration involved continuing to make the same egocentric response, as required on the response version, when the trial required turning the opposite direction to enter the visual-cue arm. For every consecutive 12 trials in a session, half the trials consisted of these trials. These trials were separated into consecutive blocks of 4 trials each (Ragozzino, 2002). Perseveration was defined as entering the incorrect arm in 3 or more trials per block. This is a similar criterion as used in previous experiments measuring perseveration (Ragozzino et al., 1999; Floresco et al., 2006). Once a rat made less than three errors in a block the first time, all subsequent errors were no longer counted as perseverative errors.

## Acknowledgments

This work was funded by FONCyT (PICT 2015-0110 to PB and PICT 2015-2344 to NW), Consejo Nacional de Investigaciones Científicas y Técnicas (PIP 0564 to PB), IBRO Return Home Fellowship to PB and a Medical Research Council grant (MC-A060-5PR00) to MCA. The authors would like to thank David Jaime for helping us with the animal maintenance.

